# Slow and not so furious: *de novo* stomatal pattern formation during plant embryogenesis

**DOI:** 10.1101/2022.09.02.506417

**Authors:** Margot E Smit, Anne Vatén, Andrea Mair, Carrie A M Northover, Dominique C Bergmann

## Abstract

The fates of cells on the leaf surface depend on positional cues and environmental signals. New stomata are formed away from existing ones to prevent disadvantageous clustering, but how are the initial precursors positioned? Mature embryos do not have stomata, but we provide transcriptomic, imaging, and genetic evidence that Arabidopsis embryos engage known stomatal fate and patterning factors to create regularly spaced stomatal precursor cells. Analysis of embryos from 35 plant species indicates this trait is conserved across angiosperm clades. Embryonic stomatal patterning in Arabidopsis is established in three stages: first broad SPEECHLESS (SPCH) expression, then coalescence of SPCH and its targets into discrete domains, and finally one round of symmetric division to create stomatal precursors. Lineage progression is then halted until after germination. We show that embryonic stomatal pattern enables quick stomatal differentiation and photosynthetic activity upon germination but also guides the formation of additional stomata as the leaf expands. In addition, key stomatal regulators are prevented from driving the fate transitions they can induce after germination, revealing stage-specific layers of regulation that control lineage progression during embryogenesis.

## Introduction

Plant and animal tissues contain distinct cell types arranged in species-specific patterns, and these patterns often facilitate optimal tissue function. The origins of pattern have fascinated developmental biologists and mathematicians for decades and engender lively debates about whether tissues require local organizers or whether they are largely self-organized. Among self-organizing paradigms, reaction-diffusion (RD) or “Turing” models capable of organizing a homogenous field of cells into discrete states dispersed across that field have been proposed for both plant and animal systems (Landge et al., 2020; Schweisguth and Corson, 2019; Xu et al., 2021).

The epidermis of Arabidopsis leaves contains trichomes and stomata, specialized cell types distributed in ways that suggest their patterns emerge from self-organizing systems (Balkunde et al., 2020; Horst et al., 2015). In the case of stomata, bi-cellular valves used for plant-atmosphere gas exchange, fate and pattern emerge through a series of state changes and oriented cell divisions (Figure 1A-B) (Lee and Bergmann, 2019). A stomatal lineage begins when a protodermal cell acquires meristemoid mother cell (MMC) identity and divides asymmetrically, creating a small meristemoid (M) and larger stomatal lineage ground cell (SLGC). Meristemoids can continue asymmetric division or transition to guard mother cell (GMC) identity before dividing symmetrically to form a pair of stomatal guard cells (GCs) that will differentiate and form a pore between them. Multiple, dispersed stomatal lineages emerge across the leaf. Each lineage may progress through precursor stages asynchronously, but cells in all lineages must communicate to form the overall pattern (Figure 1A).

**Figure 1.**
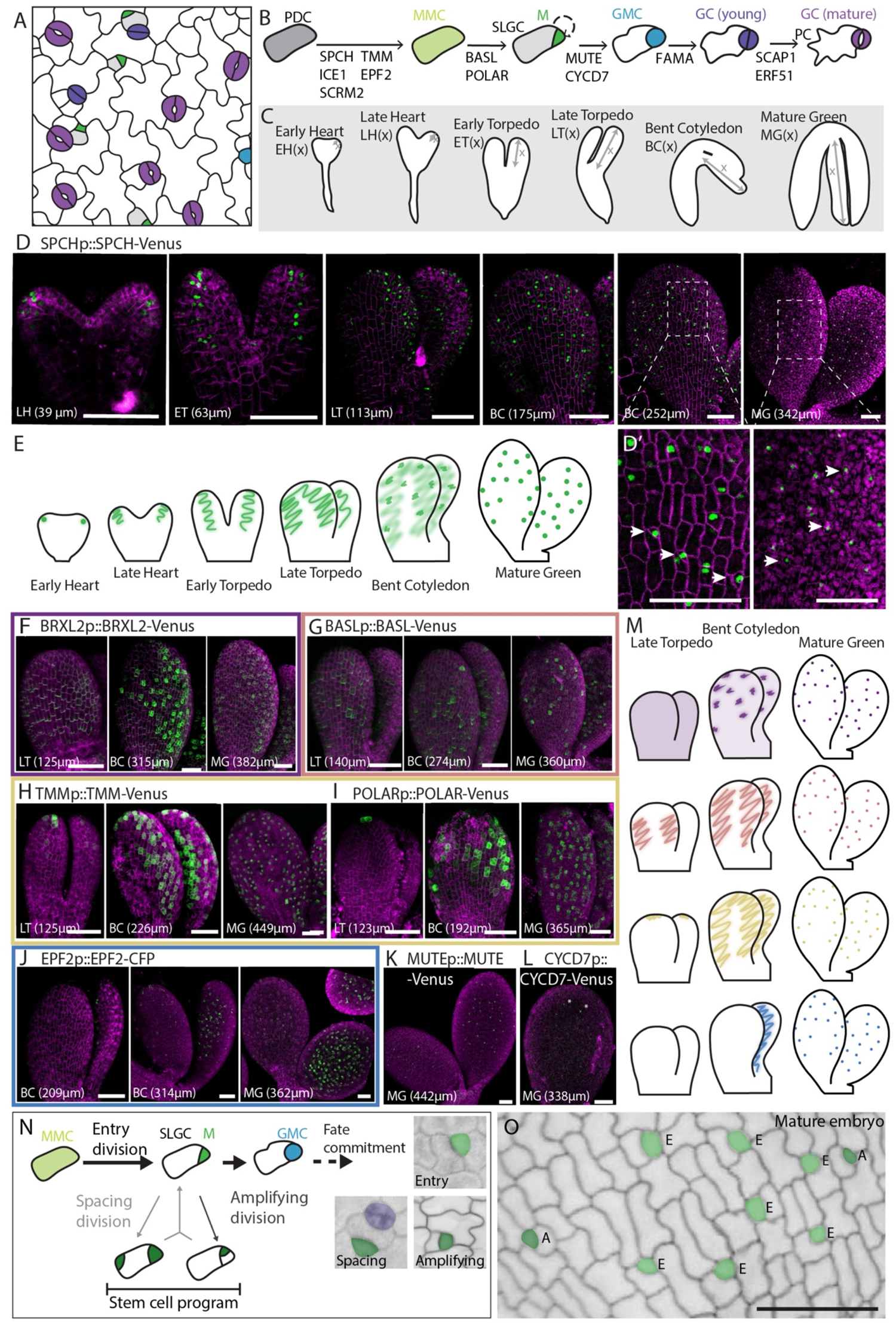
Reporters of stomatal regulators indicate presence of organized stomatal lineages in embryos. (A-B) Schematic representation of the stomata and their precursors patterned in the leaf epidermis (A) and displayed as a developmental lineage with key regulatory genes and their points of action (B), colors indicate cell identities. (C) Scheme of Arabidopsis embryogenesis annotated with stage abbreviations and region reported (x) as cotyledon length. (D) Confocal images displaying localization pattern of SPCHp::SPCH-YFP reporter (green) during embryogenesis, stages as in (C). All images are sums of stack and signal is exclusively epidermal. (D’) Lineage initiation through ACD and SPCH expression (green) in single cells, white arrows indicate small cells resulting from ACDs. (E) Graphical summary of embryonic SPCH expression in the cotyledon protoderm. (F-L) Confocal images displaying expression patterns of BRXL2 (F), BASL (G), TMM (H), POLAR (I), EPF2 (J), MUTE (K) and CYCD7 (L) translational reporters (green) during embryogenesis. (M) Graphical summary of four observed expression patterns for SPCH targets during embryogenesis. Colors are paired with micrograph outlines for reporters in F-J. (N) Overview of stem-cell like ACDs in early stomatal lineage highlighting patterns formed by Entry, Spacing and Amplifying divisions. (O) Protoderm of mature Arabidopsis embryo with LPs false colored in green, E indicates a cell resulting from an Entry division, A resulting from an Amplifying division. In D, D’ and F-L, plasma membrane marker ML1p::mCherry-RCI2A is in magenta and stomatal lineage reporters are in green. In N-O, membranes are marked by FM4-64 (N) or mPS-PI staining (O). Scale bars represent 50 μm, tags indicate embryo stage (cotyledon length). **See also Figure S1**

The identities of stomatal GCs and their precursors are guided primarily by the action of bHLH TFs, including stage-specific SPEECHLESS (SPCH), MUTE and FAMA and their more broadly expressed partners INDUCER OF CBF EXPRESSION 1/ SCREAM (ICE1/SCRM) and SCRM2 (Figure 1B) (Hachez et al., 2011; Han et al., 2018; Kanaoka et al., 2008; Lau et al., 2014; MacAlister et al., 2007; Ohashi-Ito and Bergmann, 2006; Pillitteri et al., 2007; Seo et al., 2022). The stomatal developmental pathway can be roughly divided into two phases: A SPCH-dominated proliferative stage during which overall cell number increases and the pattern emerges, and a second commitment and differentiation phase initiated by MUTE and completed by FAMA during which GMCs divide symmetrically and stomata differentiate. Ectopic expression of SPCH results in increased cell divisions and high numbers of small cells while MUTE or FAMA overexpression, or an overactive version of SCRM, induces the formation of ectopic guard cells (Kanaoka *et al*., 2008; Ohashi-Ito and Bergmann, 2006; Pillitteri *et al*., 2007).

Patterning of stomata with a “one cell spacing” rule, in that two mature stomata are never directly adjacent (Figure 1A), facilitates optimal stomatal function (Dow et al., 2014; Franks and Farquhar, 2007). Patterning involves orienting cell divisions and restricting the fate of precursors, activities that require cell-cell signaling and cell polarity programs that are both induced by, and feedback on, SPCH (Herrmann and Torii, 2021; Lau *et al*., 2014). Mobile signaling peptides in the EPIDERMAL PATTERNING FACTOR family expressed from meristemoids and GMCs engage receptors ERECTA (ER) and TOO MANY MOUTHS (TMM) to induce MAPK signaling and subsequent SPCH phosphorylation and degradation (Bhave et al., 2009; Hara et al., 2007; Hunt and Gray, 2009; Lin et al., 2017; Yang and Sack, 1995). BREAKING OF ASYMMETRY IN THE STOMATAL LINEAGE (BASL), BREVIS RADIX-LIKE 2 (BRXL2) and POLAR LOCALIZATION DURING ASYMMETRIC DIVISION AND REDISTRIBUTION (POLAR) polarly localize and are inherited by SLGCs. These “polarity proteins” scaffold kinases that ultimately result in SPCH phosphorylation and degradation (Dong et al., 2009; Houbaert et al., 2018; Rowe et al., 2019; Zhang et al., 2015). Consequently, SPCH-expressing cells induce transcription of proteins that serve to downregulate SPCH in surrounding cells, creating zones with reduced ‘stomatal competence’

The current view of stomatal development in leaves requires no pre-existing information to bias the placement of stomata and relies entirely on cell communication and error correction (Geisler et al., 2000; Horst *et al*., 2015). This is consistent with the observation that mature embryos do not have stomata (Esau, 1953; Peterson and Torii, 2012). Mature embryos from Arabidopsis and soybean, however, have epidermal cells that resemble meristemoids, suggesting that the stomatal lineage is initiated during embryogenesis (Danzer et al., 2015; Geisler and Sack, 2002). Moreover, expression profiles of soybean revealed embryonic transcripts of *GmSPCH*, and their depletion eliminated the embryonic meristemoid-like cells (Danzer *et al*., 2015).

Armed with contemporary transcriptomic data, imaging innovations to survey plant embryos broadly, and tools to monitor and manipulate embryonic gene expression in Arabidopsis, we revisit the question of whether stomatal precursors are stochastically chosen in cotyledons, or whether they reflect an embryonic pre-pattern. We track the stages of initial stomatal pattern formation using a set of stomatal translational reporters and find that the early stomatal lineage is multiphasic, with a delay between SPCH and SCRM accumulation and their full activity, as marked by target gene expression and asymmetric cell divisions. A phenotypic analysis of 35 angiosperm species demonstrates that embryonic stomatal lineage activities are frequent and widely distributed. Genetic manipulations in Arabidopsis allowed us to show that embryonic stomatal patterning enables rapid stomatal differentiation and leaf expansion upon germination and facilitates efficient patterning of stomatal lineage expansion in seedlings. By changing the expression and activity of SPCH, SCRM, MUTE and FAMA in embryos, we show how developmental stage intersects with transcriptional activity, showing, for example, that precocious MUTE or FAMA expression alters cell morphology, division, and gene expression but cannot induce guard cell identity during embryogenesis. Together these findings highlight the power of comparing embryonic and post-embryonic elaboration of developmental programs to uncover patterning rules and to refine the gene regulatory networks (GRNs) surrounding key developmental transcription factors.

## Results

Although plant embryos are widely reported to lack stomata, two recent Arabidopsis transcriptional profiles revealed extensive stomatal development gene expression in embryos (Figure S1A-D)(Hofmann et al., 2019; Schneider et al., 2016), and a paper characterizing embryonic cotyledon growth noted expression of reporters for stomatal lineage receptor genes (Chen and Shpak, 2014). We wondered whether gene expression indicates an organized developmental program. If it does, how extensive is this stomatal program, and what is its purpose, if not to create stomata? The temporal patterns of stomatal gene expression from the embryo transcriptome experiments largely recapitulate the patterns in leaves, but transcriptomes lack information about spatial distribution and gene function. Thus, we set out on a systematic characterization of stomatal gene reporter behavior and of the effects of embryonic loss or gain of stomatal gene function. We envisioned several functions for stomatal genes in embryos: (1) activity of the stomatal lineage in embryos could create cells primed to differentiate into stomata immediately upon germination to provide a baselines of gas exchange, (2) “pre-patterned” cells could facilitate a more orderly emergence of stomatal pattern upon germination, and/or (3) because stomatal lineages produce the majority of epidermal cells, including non-stomatal cells, in leaves (Geisler *et al*., 2000) stomatal genes may be recruited for general cell proliferation-promoting activities.

### Embryonic expression of SPCH starts early and wide before becoming restricted to stomatal lineage cells

We began by characterizing the pattern of a functional SPCH translational reporter (SPCHp::SPCH-YFP in *spch*)(Lopez-Anido et al., 2021) because SPCH, with its partner SCRM, marks stomatal lineage initiation. We detected SPCH as early as heart stage in the cotyledon protoderm (Figure 1D-E). For simplicity, we will refer to the outer tissue layer as protoderm when describing the embryo and epidermis when describing the germinated seedling. Additionally, we classify early stomatal lineage cells based on multiple stereotyped criteria (see methods) and will refer to cells from lineage entry until just before the final symmetric division to form guard cells as “Lineage Precursors (LPs)” in embryos.

During initial cotyledon expansion until the early torpedo stage, SPCH protein is uniformly present in protodermal cells at the leaf edge (Figure 1D-E, S1E). A reporter for SPCH’s partner SCRM (SCRMp::SCRM-mCit) can be detected broadly in protodermal and internal tissues from early embryo development onwards (Figure S1F). The SPCH expression domain narrows as embryogenesis progresses and by the mature green stage, SPCH is exclusively present in small LPs representing the smaller daughter cells of ACDs. Revisiting SPCH expression in true leaves, we find it initially present in a broad band of cells at the edge of the first true leaves (Figure S1H). Additionally, in the Brachypodium embryo, BdSPCH2 is expressed in the protoderm of the first leaf from leaf early stage onwards (Figure S1J)(Hao et al., 2021; Raissig et al., 2016). Thus, initially broad SPCH presence is common during leaf initiation despite vastly different morphologies exhibited by monocots and dicots.

### Stomatal signaling and polarity factors coalesce into patterns resembling SPCH by bent cotyledon stage, but display diverse initial patterns

Translational reporters of representative signaling (EPF2, TMM) and polarity (BASL, POLAR and BRXL2) factors previously demonstrated to be SPCH targets could be detected in patterns associated with SPCH from the late torpedo stage onward (Figure 1F-J). Interestingly, this shared pattern late in embryogenesis represents a coalescence of expression from initially diverse sources. Uniquely, BRXL2 precedes SPCH and is initially present in all protodermal cells where it appears to report a global polarity field (Figure 1F). Only later, at bent cotyledon stage, does BRXL2 increase in LPs and fade from other cells. BRLX2 remains polarized in LPs but is no longer globally oriented towards the organ base. Polarity protein BASL is first observed at the base and middle of early torpedo stage cotyledons in small groups of cells before it associates with the SLCGs after ACDs (Figure 1G, S1K). TMM and POLAR most closely resemble SPCH itself in their expression: first present in the tip of the cotyledon at the late torpedo stage before expanding and finally gradually narrowing to small domains of expression (Figure 1H-I, S1K). Finally, the signaling protein EPF2 is detectable later than all other SPCH targets, starting at bent cotyledon stage, (Figure 1J, S1K). EPF2 expression also coincides with the appearance of ACDs, a hallmark of stomatal lineage activity (Figure 1D-D’). Interestingly, based on cell shapes, neighbor orientations and overall cell pattern before germination, embryonic ACDs are primarily entry divisions (Figure 1N-O); embryonic SLGCs do not undergo spacing divisions and meristemoids rarely undergo amplifying divisions.

Finally, as embryos reach maturity, we find MUTE expressed in many of the LPs (Figure 1K, S1K) and CYCD7;1 expressed sporadically and at very low levels (Figure 1L). In contrast, FAMA and downstream stomatal differentiation factors, including SCAP1 and ERF51, are not detected during embryogenesis, neither as transcripts (Figure S1C-D), nor as transcriptional or translational reporters (negative data not shown).

### Mutations in stomatal lineage genes affect patterning of the embryonic protoderm

The convergence of expression of many stomatal regulators in the embryonic protoderm prompted us to test whether these stomatal factors are required for the embryonic stomatal pattern. Based on the phenotypes of *SPCH* loss and gain of function mutants in Arabidopsis leaves and the lack of LPs in *GmSPCH* RNAi embryos (Danzer *et al*., 2015; MacAlister *et al*., 2007), we expected to see changes in ACD frequency and in numbers of LPs when we altered *SPCH* or *SCRM*. Because embryos do not have stomata, we expected the loss of *MUTE* or *FAMA* to have negligible effects on embryos. More interesting was the question of whether *EPF2, TMM, BASL, BRX-q*, genes implicated in preventing mature stomata from being formed in contact and/or enforcing division and fate asymmetry, would have any role in the protoderm that appears to undergo ACDs, but lacks stomata.

We characterized the effects of mutations in stomatal development genes on the protodermal patterns of mature embryos by dissecting imbibed seeds, staining with mPS-PI and extracting surface signal from processed confocal stacks with MorphographX (details in methods). SPCH levels correlate with protodermal cell numbers, with mature *spch* embryos lacking small LPs (Figure 2A-B) and an overexpression line (ML1p::SPCH) having more (Figure 2C). Embryos expressing a hyperactive *SCRM-D* allele similarly have increased numbers of small LPs (Figure 2D). Strikingly, in contrast to its postembryonic phenotype (Kanaoka *et al*., 2008), *SCRM-D* did not produce ectopic guard cells in embryos. Coupled with the lack of discernable protodermal phenotypes in *mute* embryos (Figure 2E), it appears that that lineage progression past the GMC stage is not possible during embryonic development.

**Figure 2.**
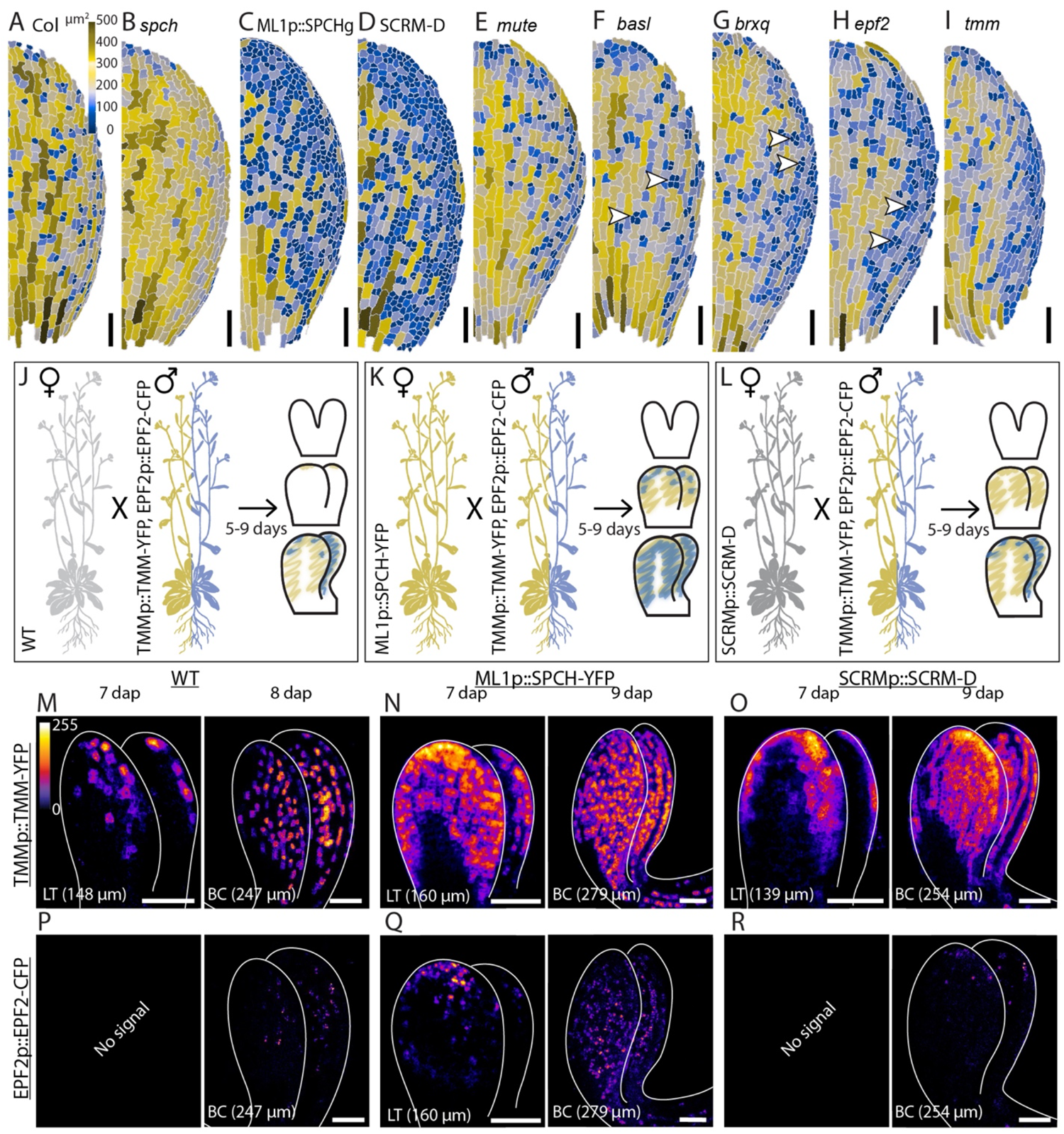
Embryonic phenotypes resulting from loss or gain of function mutations in stomatal regulators. (A-H) Morphographx-segmented mature embryos of Col-0 (WT) (A) *spch* (B), ML1p::SPCH (C), SCRM-D (D), *mute* (E), *basl* (F), *brxq* (G), *epf2* (H) and *tmm* (I); color of segmented cells indicates cell area with key in (A). Arrowheads point to examples of clustered LPs. (J-L) Schematic representation of experimental setup to test effects of SCRM-D or increased SPCH on TMM and EPF2 at 7-9 dap. (M-R) Representative confocal images of TMM (M-O) and EPF2 (P-R) expression in WT (M,P), ML1p::SPCH-YFP (N,Q) and SCRM-D (O,R) embryos. Fluorescent signal intensity on Fire LUT (M-R). Scale bars represent 50 μm, tags indicate Embryo Stage (cotyledon length). **See also Figure S2**

Patterning mutants *basl, brxq* and *epf2* exhibited a common phenotype of clustered small LPs at the end of embryogenesis, a phenotype paralleling their seedling roles, and suggesting that even without stomata, lineage precursors employ signaling and polarity mechanisms (Figure 2F-H). Loss of *TMM*, unexpectedly, resulted in fewer LPs (Figure 2I), resembling *tmm*’s stem rather than its leaf phenotypes (Bhave *et al*., 2009; Yang and Sack, 1995), hinting that signaling may underlie differences in embryonic and postembryonic patterning rules.

### Precocious and higher levels of SPCH or SCRM-D cannot fast-track embryonic stomatal development

Our expression and functional data indicate that components of the early stomatal lineage are active in embryos, but we also see a lag between early expression of SPCH and SCRM, the activation of their target genes, and the appearance of ACDs. To test whether this lag, which is not seen in seedlings, was due to insufficient levels of SPCH and SCRM, or due to a stage-specific lack of competence to respond to these regulators, we crossed lines expressing ML1p::SPCH or SCRM-D with lines bearing translational reporters for TMM and EPF2 (Figure 2J-L). After crossing, embryos were collected and imaged 5-9 days after pollination (dap) to capture late heart to bent cotyledon stage embryos.

From late torpedo stage onwards, ML1p::SPCH and SCRM-D were both able to induce additional protodermal divisions, excess LPs and high, broad TMM expression (Figure 2M-O, S2A-C). In addition, ML1p::SPCH was able to induce broader, higher, and earlier expression of EPF2 (Figure 2Q), suggesting that SPCH levels are limiting at this stage. In contrast, SCRM-D had no effect on EPF2 expression (Figure 2R). Neither ML1p::SPCH nor SCRM-D was able to induce TMM or EPF2 expression or extra divisions before late torpedo stage, however, suggesting that protodermal cells in the pre-torpedo stage embryo are not yet competent to respond to SPCH/SCRM and/or to initiate the stomatal lineage.

### Embryonic stomatal patterning is widely conserved across eudicots

Both Arabidopsis and soybean (Danzer *et al*., 2015; Geisler and Sack, 2002) have LPs at the end of embryogenesis, suggesting that initiation, but not completion of the stomatal lineage, could be a common feature in angiosperm embryos. When we imaged mature tomato embryos, however, we found no such LPs (Figure 3A, S3B). We therefore performed a broader survey of embryos to discern phylogenetic or ontogenetic patterns to the appearance of embryonic stomatal precursors. We dissected embryos from seeds of 35 eudicot and 1 monocot species (See KeyResources table) and found that small, regularly spaced protodermal cells, which resemble meristemoids or GMCs are present in several rosid and asterid species, and the one monocot we measured (Figure 3B-D and Figure S3B).

**Figure 3.**
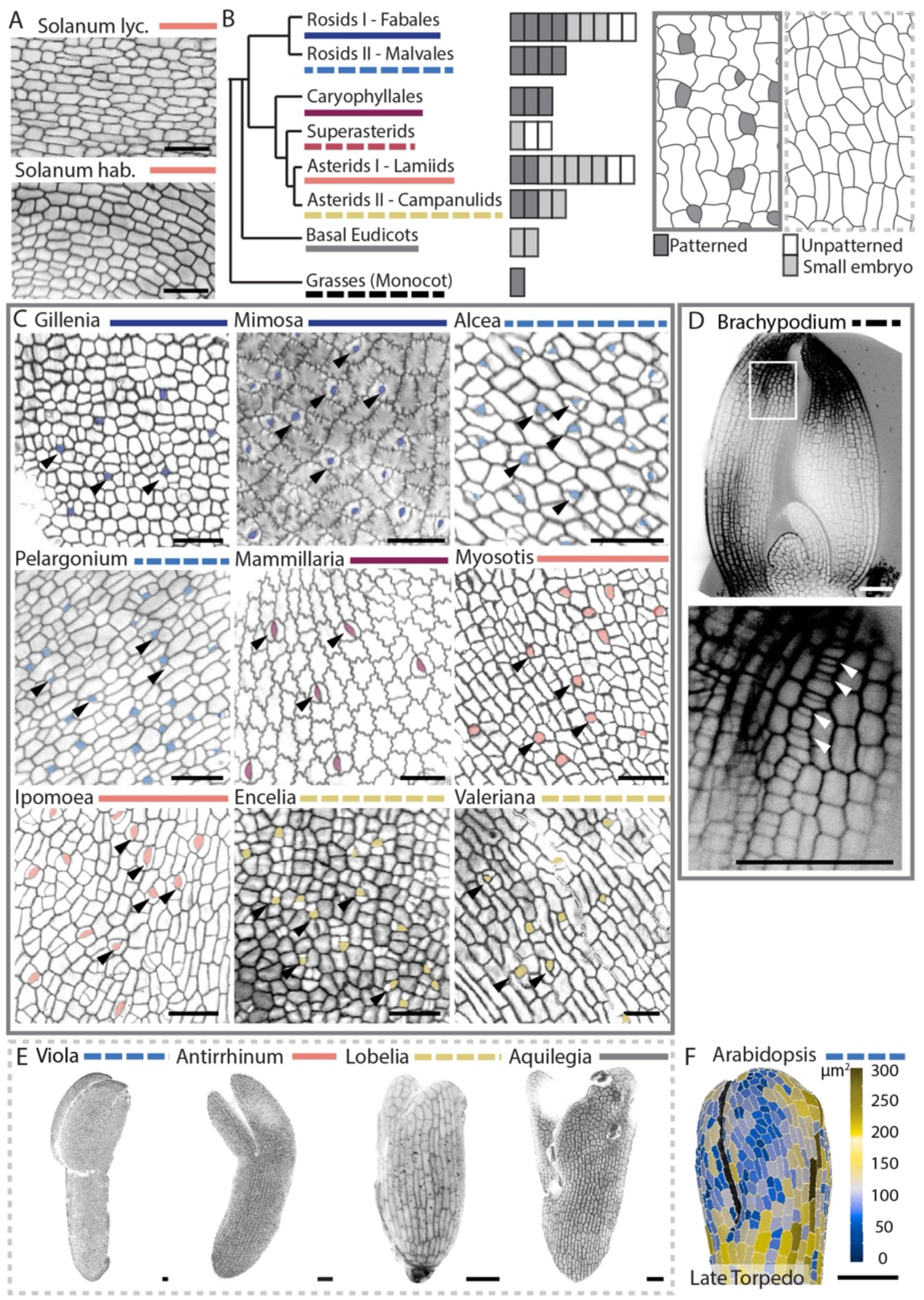
Diverse angiosperm species display evidence of stomatal lineage activities in embryos. (A) Confocal images of mPS-PI stained protodermal pattern in mature embryos of *Solanum lycopersicum* and *Solanum habrochaites*. Seeds were dissected and embryos stained with mPS-PI, black signal shows cell walls. After imaging, epidermal signal was isolated by creating a mesh and projecting epidermal signal using MorphographX (see methods). (B) Left: Cladogram showing major eudicot clades sampled in this study. Middle: Bars representing different species, dark gray bars represent species with patterned embryos, white bars indicate absence of detectable embryonic stomatal patterning, grey bars refer to species with smaller, immature embryos. Right: Drawing of a patterned vs unpatterned protoderm. (C) Protodermal pattern of several eudicot species, stained and processed as in (A). LPs are false colored and indicated with arrowheads. Colored bars represent the clade (from B) to which each species belongs. (D) LPs in embryonic leaves of *Brachypodium distachyon*, LPs are marked with arrowheads. (E) Embryo shape and epidermal patterning of eudicot species with small embryos in the mature seed. Colored bars represent the clade (from B) to which each species belongs. (F) Shape and epidermal pattern of an *Arabidopsis thaliana* late torpedo stage embryo. Color of segmented cells indicates cell area. Scale bars represent 50 μm. **See also Figure S3**

Within each clade, several plant species appeared to lack embryonic stomatal lineages. A common feature of the species without embryonic LPs is that their embryos are immature at desiccation (13/20 species). Here we define an immature embryo as taking up less than 90% of the seed volume. The shape of immature embryos can resemble earlier stages of Arabidopsis embryogenesis (Figure 3E-F). Indeed, these species (or close relatives) were described as having small embryos relative to their overall seed size and were hypothesized to continue development after desiccation and seed dispersal (Forbis et al., 2002; Lubbock, 1892; Martin, 1946). Thus, exploring natural phenotypic diversity leads to the same conclusion as a genetic and molecular dissection of Arabidopsis—that there are two phases of protodermal development, an early phase where proliferation is uncoupled from the stomatal lineage and a later stage that includes full engagement of ACDs and stomatal lineage fate, patterning, and polarity factors.

### Pre-patterned stomatal cells quickly differentiate upon germination

If embryonic initiation of the stomatal lineage is widespread, what is its function? Two hypotheses suggested by classic developmental and physiological studies (Geisler *et al*., 2000; Zeiger and Field, 1982) are that primed stomatal precursors enable quick differentiation to support gas exchange immediately after germination and/or that an initial pattern laid down in the embryo can guide ACDs and new lineage entries in a more organized and efficient manner in the young cotyledon.

To test the first hypothesis, we followed the fate of embryonically patterned stomatal lineage cells after germination. We generated germination time-courses, capturing images from individuals at radicle protrusion (24-36 hours after placement in light, here labeled as 0hr) and 8, 24 and 48 hours later (Figure 4A-D, S4A-D). Nearly all (92%, n = 159/172 cells across 3 leaves) of stomatal lineage cells identified at radicle protrusion had formed stomata within 48 hours, with the majority (63% n = 108/172 cells) immediately dividing symmetrically and forming a single pair of guard cells (Figure 4A-E). Therefore, nearly all stomatal cells formed during embryogenesis are GMCs or are a single division away from becoming GMCs. These cells indeed quickly have MUTE and FAMA expression 0-12 hours after radicle protrusion (Figure 4G-H, S4E-H) and are responsible for most stomata present 48 hours after radicle protrusion (Figure 4E). This immediate differentiation of embryonically patterned cells into stomata was also observed in *Ruta graveolens* and *Myosotis alpestris*, two other species whose seedlings were amenable to time course analysis (Figure S4I-L).

**Figure 4.**
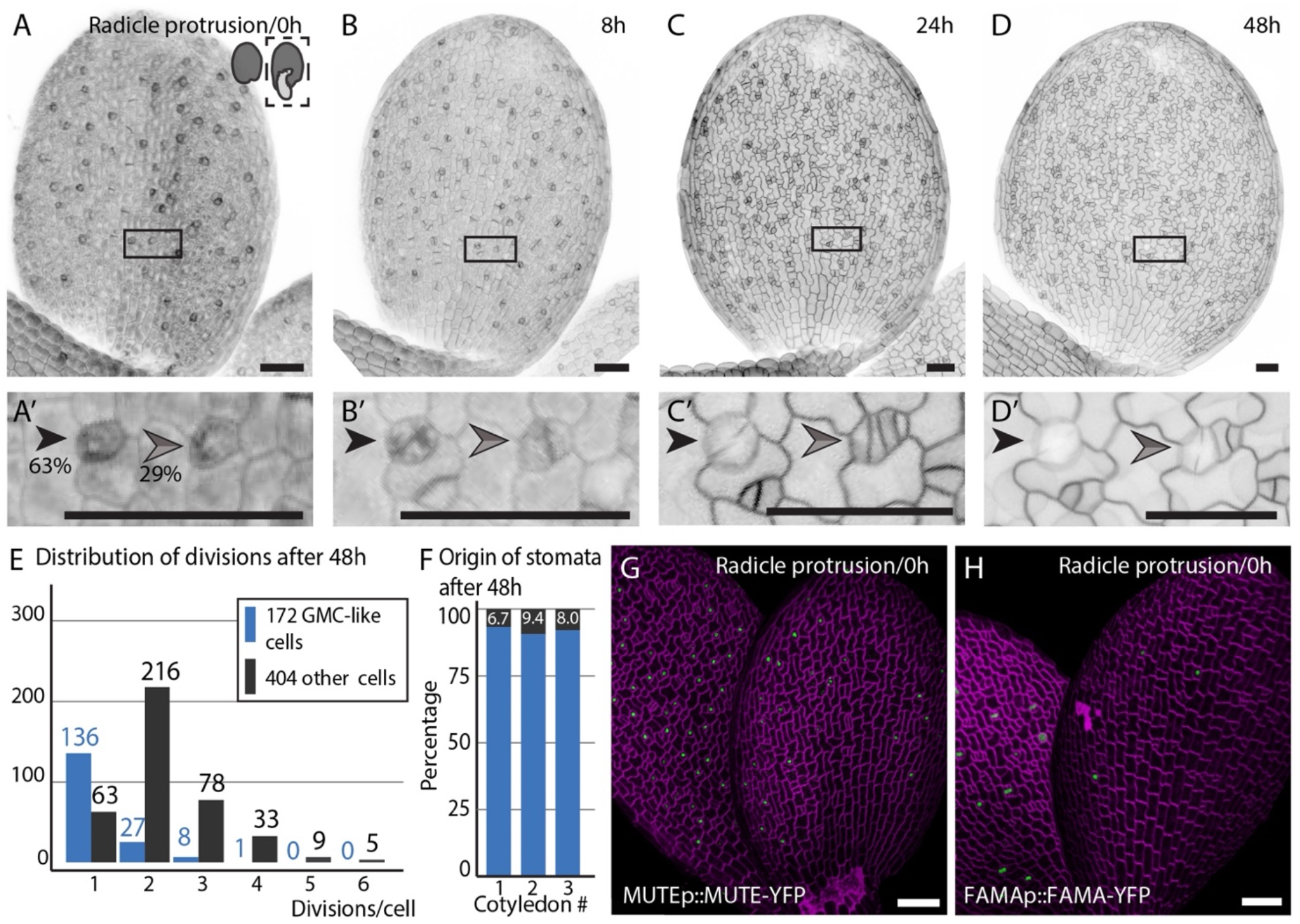
Lineage Precursors rapidly give rise to stomata upon germination. (A-D) Time course series showing epidermal cell division and differentiation 0h (A), 8h (B), 24h (C) and 48h (D) following radicle protrusion. Inserts (‘) follow two representative LPs, one directly differentiating (black arrow) and one first undergoing an extra division (grey arrow). (E) Distribution of divisions each cell has undergone in the first 48h after radicle protrusion, GMC-like cells (blue) and other dividing cells (black). (F) Bar graph showing the contribution of embryonically patterned stomatal cells (blue) and new lineage entries (black, percentage) to the total number of stomata 48 hours after radicle protrusion. (G-H) Expression of MUTE (G) and FAMA (H) translational reporters (green) at radicle protrusion. Black or magenta signal shows cell shape marked with either FM4-64 (A,B) or ML1p::mCherry-RCI2A (C-D,G-H). Scale bars represent 50 μm. **See also Figure S4**

Embryonic activities may enable the formation of stomata quickly upon germination, but how essential is this for plant growth? We tested the consequence of making stomatal precursors, but arresting their development into stomata, by following the growth of *fama* mutants. Loss of *FAMA* does not lead to detectable embryonic phenotypes (Figure S4O-P) but we found it has reduced cotyledon expansion starting 2 days after radicle protrusion (dar) (Figure S4Q). Since embryonically defined stomata make up more than 90% of stomata present at this time point, “pre-patterned” stomata appear to allow for a smooth transition to photosynthetic autotrophy.

### Without the embryonic stomatal lineage, post-embryonic lineage entry is disorganized, resulting in faulty epidermal patterning

Next, we investigated whether embryonic stomatal lineage cells contribute to post-embryonic stomatal patterning. We created an inducible *spch* rescue line (Figure 5A) to test the effects of inducing stomatal development post-embryonically in the absence of any embryonic SPCH (and by extension, in the absence of ACDs and SPCH target genes involved in cell communication and polarity). If SPCH activity is required for *stomatal identity competence*, then we would expect few post-embryonic stomatal lineage divisions upon SPCH induction. Alternatively, if embryonic activity serves a patternin*g* role, i.e., to restrict “stomatal neighbor” cells from entering the lineage, then excessive and clustered precursor cells would appear even at low levels of SPCH induction.

**Figure 5.**
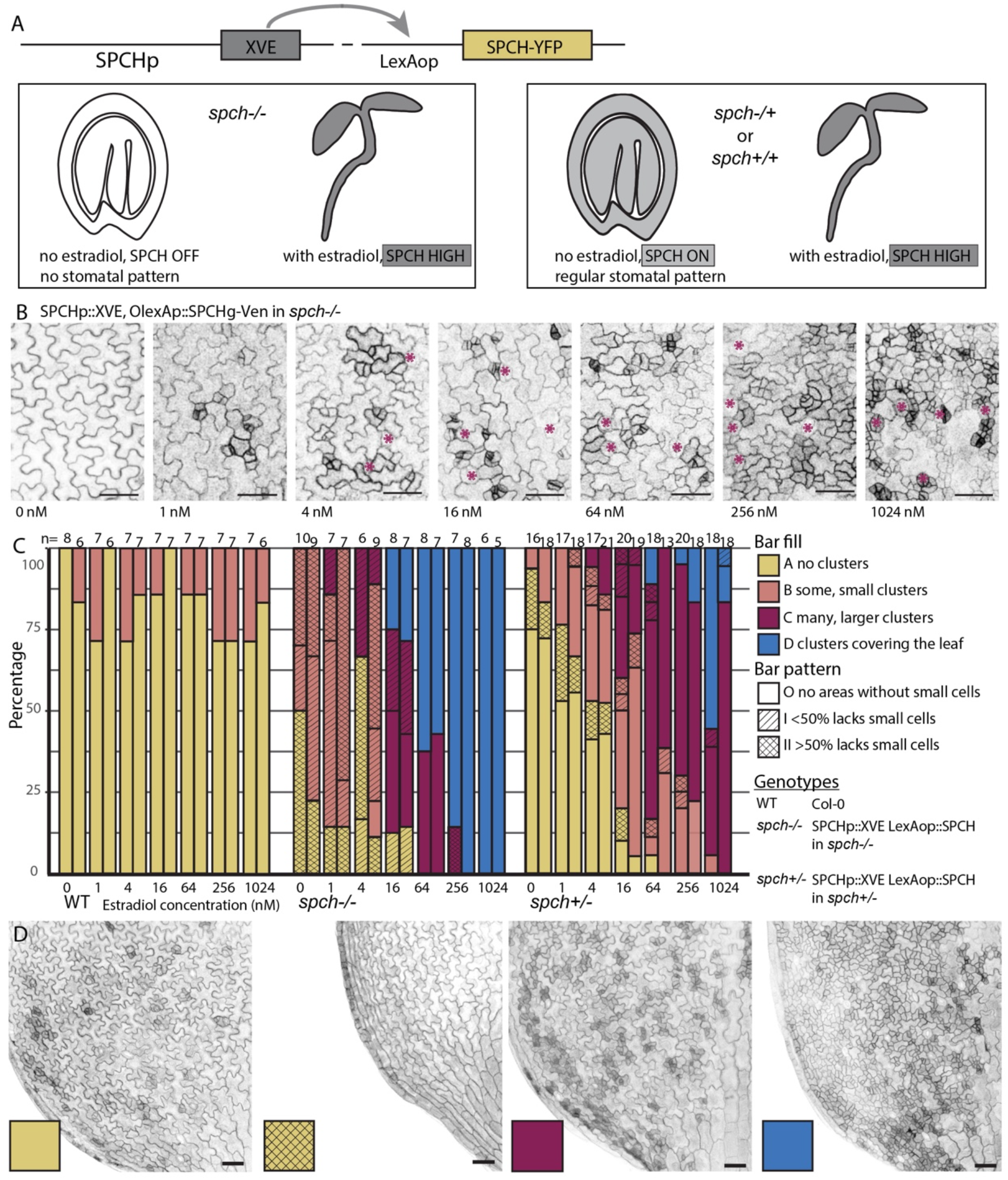
Embryonic stomatal lineage activity guides post-embryonic stomatal patterning. (A) Schematic overview of experimental setup for inducible SPCH and levels of SPCH (shades of grey) that different genotypes experience. (B) Confocal images of 4-dag *spch* cotyledon epidermis, showing increase in stomata and stomatal precursors with increasing estradiol concentrations. Purple stars mark differentiated stomata. (C) Summary of stomatal identity and pattern phenotypes in 4 dap cotyledons across 3 genotypes and 7 estradiol concentrations. X-axis shows 3 genotypes (WT, SPCHp>>SPCH in *spch-/-*, SPCHp>>SPCH in *spch+/-*) and 7 estradiol concentrations tested per genotype. Two replicates, each measuring 6-21 leaves, n indicated at top are displayed for each timepoint or genotype. Cotyledon phenotypes are shown by a combination of bar fill color (small cell phenotype) and bar pattern (fraction of area without small cells or stomata). (D) Confocal images of representatives of phenotypic classes (indicted by colored and patterned boxes) quantified in (C). Black signal marks cell outlines using ML1p::mCherry-RCI2A. Scale bars represent 50 μm. **See also Figure S5**

As expected, without SPCH induction, 4-dag *spch* seedlings created no stomata (Figure 5B). Increasing levels of post-embryonic SPCH induction resulted in additional cell divisions but none of the seven estradiol concentrations used (0, 1, 4, 16, 64, 256, and 1024 nM) resulted in a regularly patterned epidermis: either too few stomata were formed alongside clusters of small cells, or small cell clusters dominated the leaf surface (Figure 5B,D). This suggests embryonic stomatal lineages serve a patterning role and, in their absence, postembryonic lineage entry is disorganized and excessive. One caveat to our experimental set-up was that it was possible that the disorganized post-embryonic development was not due to missing patterning cues, but rather the result of excessive SPCH induction. We therefore confirmed that *spch* seedlings did not accumulate more SPCH than *spch+/-* or *SPCH+/+* individuals at low induction levels (Figure S4A-C) and repeated the induction experiments in a *spch+/-* heterozygous line (Figure 5A,C). At high levels of induction (256 or 1024 nM), ectopic divisions are found in both lines, as expected. However, at low to medium levels of induction (4 – 64 nM) the clustering phenotypes in the *spch+/-* population are less severe than in *spch-/-* individuals (Figure 5C-D), indicating that embryonic stomatal lineage patterning makes cotyledons less likely to experience excessive and mispatterned lineage entry after embryogenesis.

### Embryonic misexpression of MUTE or FAMA does not result in the formation of stomata

Morphological and molecular markers indicate that embryonic stomatal development halts at the GMC stage; we never observe FAMA transcriptional or translation reporter expression. If progression to stomatal identity is controlled by the transcriptional regulation FAMA, then we should be able circumvent the arrest by artificially expressing FAMA in embryos. We first expressed FAMA under the MUTE promoter but could not induce stomata during embryogenesis (Figure 6D-E). The same MUTEp::FAMA-YFP line, generates single, kidney-shaped guard cells after germination (Figure 6A), however, indicating that it is the embryonic context, not the transgene that defines the ability to produce stomata.

**Figure 6.**
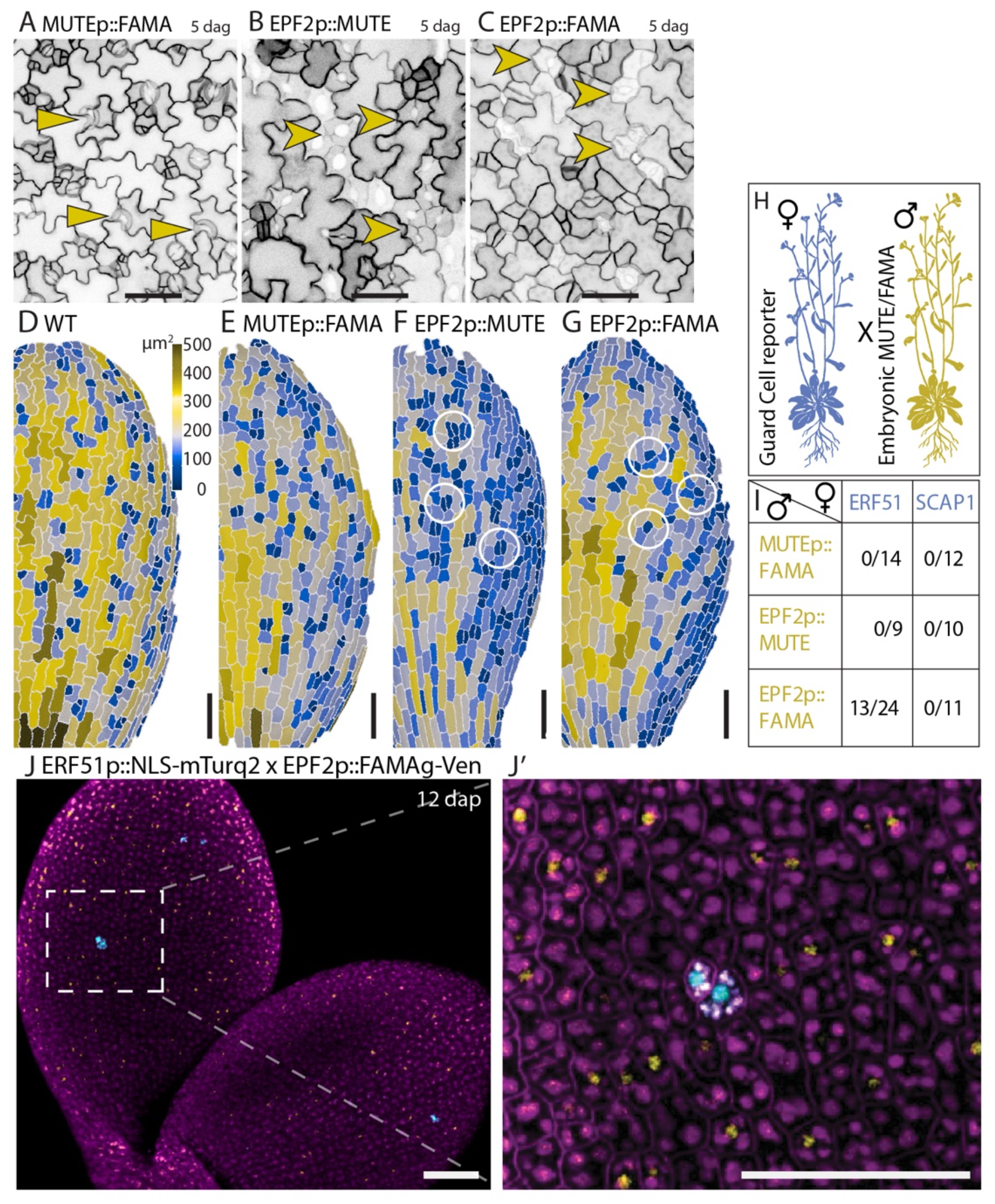
Misexpression of MUTE or FAMA during embryogenesis cannot trigger stomatal guard cell formation. (A-C) Confocal images of 5-dag cotyledon epidermis showing that precocious FAMA (A,C) or MUTE (B) expression creates ectopic GCs post-germination; arrows indicate single guard cells (A) and ectopic guard cell pairs (B,C) 5 dag. (D-G) MorphographX-segmented images of mature embryos showing that before germination, precocious FAMA (E, G) or MUTE (F) expression results in abnormal cells or extra symmetric divisions (indicated with circles) but no guard cells. (H) Schematic representation of crosses between lines precociously expressed MUTE or FAMA lines and guard cell reporter genes. (I) Table: Number of embryos 12 dap from each cross with ERF51/SCAP1 expression. (J) Confocal image of sporadic ERF51 expression (cyan) activated by EPF2p::FAMAg-Venus misexpression 12 dap. Black or magenta signal ML1p::mCherry-RCI2A (A-C,J) Blue/yellow color indicates cell area (D-G). Yellow/cyan signal a stomatal reporter/misexpression (I). Scale bars represent 50 μm.

The block at the end of embryogenesis is before the final division that creates guard cells, so we also tested whether earlier and higher expression of MUTE or FAMA could generate embryonic stomata. We used the EPF2 promoter, as we had shown that EPF2 is expressed later than other SPCH targets but well before, and at higher levels than MUTE, in embryos (Figure 1J, S1A-B). After germination, expression of either MUTE or FAMA under the EPF2 promoter resulted in the formation of ectopic and oddly shaped stomatal complexes (Figure 6B-C). Again, however, during embryogenesis no guard cells were formed despite evidence that FAMA and MUTE translational reporters do accumulate (Figure 6F-G,J). Interestingly, ectopic MUTE could induce extra symmetric cell divisions and FAMA induced cell shape changes, developing slightly elongated GMCs in the mature embryo (Figure 6F-G).

While the abnormal cells in mature embryos of MUTE and FAMA misexpression were not guard-cell shaped, the same can be said for many of the abnormal pore-forming cells in leaves produced by these lines (Figure 6B-C). Thus, we considered the possibility that MUTE and FAMA could direct transcription of some, but not all, of their downstream targets, and monitored expression of guard cell reporters in EPF2p::MUTE-YFP and EPF2p::FAMA-YFP embryos 12 days after pollination (Figure 6H). MUTE was not able to induce expression of either reporter, but FAMA could induce sporadic expression of ERF51, a direct target (Hachez *et al*., 2011)(Figure 6I-J). Taken together, we conclude that MUTE and FAMA can execute portions of their regulatory repertoire during embryogenesis, including expression of target genes, promoting symmetric divisions and inducing cell shape changes, but neither TF is sufficient to induce stomatal guard cell identity and differentiation programs.

## Discussion

We find that in the Arabidopsis embryo, stomatal lineage gene activities are highly organized in time and space, with several developmental blocks slowing down stomatal lineage progression. The outcome of this organization is a protoderm with cells primed to become stomata and other cells pre-patterned to resist later SPCH activity and thereby facilitate the orderly elaboration of the stomatal pattern upon germination.

### Hard and soft developmental blocks limit stomatal TF functionality during embryogenesis

The effects of SPCH, SCRM, MUTE and FAMA expression have been characterized in some detail in leaves, where each is sufficient to initiate major cell state changes. In the embryo, however, we find that their capacity to induce divisions, morphological change and gene expression is limited. For SCRM, MUTE and FAMA, this manifests itself as a failure to promote mature stomatal guard cell fates. Earlier in the lineage, SPCH expression and activity reveal distinct spatial and temporal phases slowing down lineage entry through ACD.

We find that expression of SPCH target genes significantly lags behind SPCH, and among targets find considerable diversity in initial expression timing and location across the cotyledon (Figure 7A-B). In general, transcription factors have unique targets in different cell types or organs, resulting from transcription factor expression level, cofactor availability, or signaling from other tissue layers (Johnson et al., 2006; Loker et al., 2021; Painter et al., 2018; Schweisguth and Corson, 2019). Examples include SHORTROOT regulating unique genes in the root cortex, xylem and quiescent center (Cui et al., 2011) and Ultrabithorax affecting gene expression in the Drosophila wing and haltere (Loker *et al*., 2021). We see instead see heterogeneity in SPCH target expression across the same tissue and organ. This heterogeneity in SPCH activity does not appear to be due to heterogeneity in SPCH levels per cell, as SPCH is initially fairly uniform across the protoderm (Figure 1D), although increasing SPCH protein levels overall can force “late” targets earlier. In the seedling, SPCH’s dimerization partners, SCRM and SCRM2 both appear to compensate for the other’s absence (Kanaoka *et al*., 2008). However, we find that they have different expression patterns in the embryo, suggesting that SPCH/SCRM and SPCH/SCRM2 heterodimers could activate different targets during embryogenesis (Figure S1F-G).

**Figure 7.**
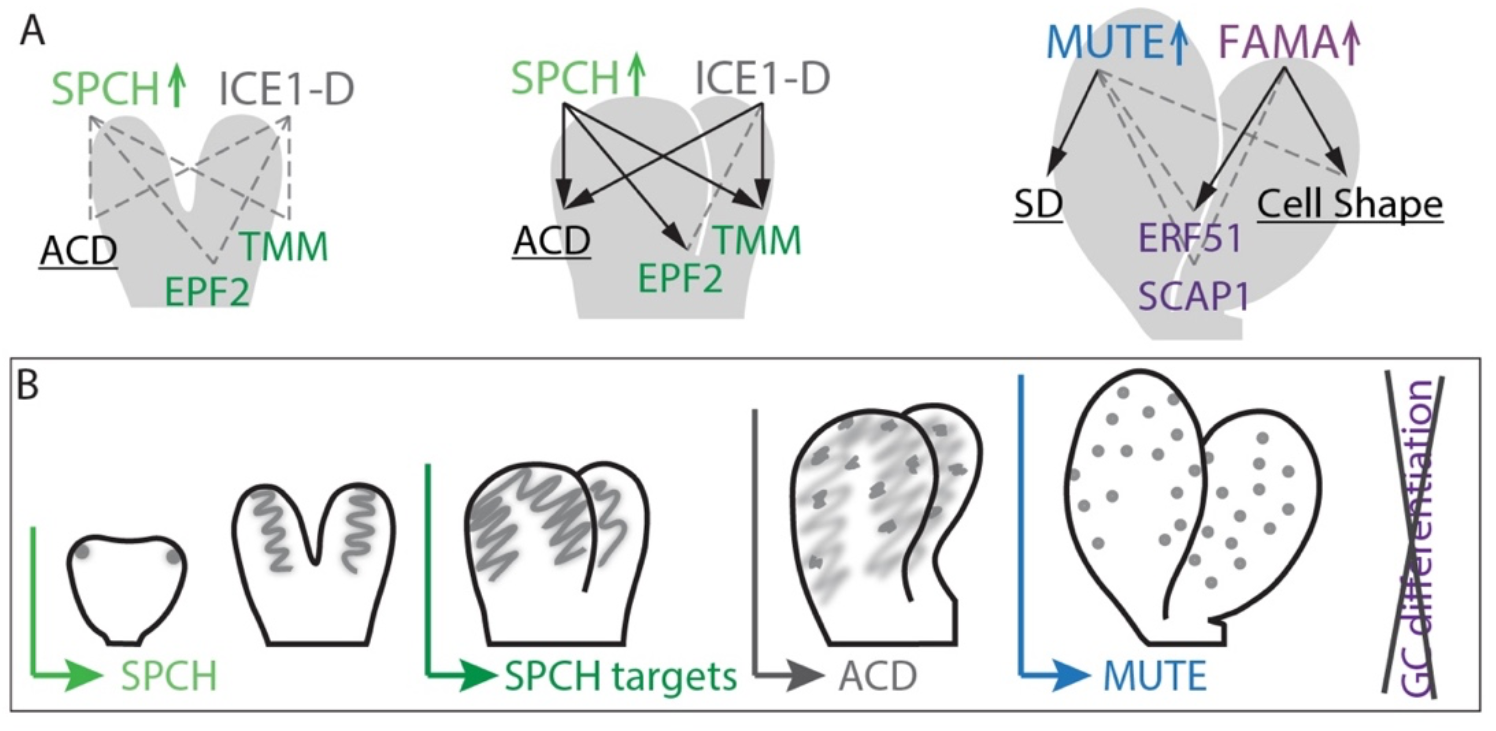
Action of stomatal transcription factors during embryogenesis. (A) Model of effects of SPCH/SCRM-D/MUTE/FAMA misexpression during embryogenesis. Schematic overview of known effects of misexpression after germination, missing effects at a particular embryonic stage are shown with dashed, grey line. In summary, SPCH and SCRM-D can induce ACD and target gene expression at torpedo stage but not earlier (left, middle), MUTE and FAMA can only alter development but cannot induce the formation of guard cells (right). (B) Timeline displaying key events and transitions during embryonic stomatal development. Here we find that some of these events can be induced earlier (ACD induction, EPF2 expression) while others are prevented at earlier developmental stages (GC differentiation, SPCH target gene expression before late torpedo stage).

Even after all monitored SPCH targets have appeared, SPCH-induced ACDs are further delayed until the final stages of embryogenesis. The late initiation of ACDs appear to be a conserved feature as we find LPs to be absent in species with relatively “young” embryos at seed desiccation (Figure 3E). In Arabidopsis, SPCH target expression and ACDs can be forced earlier through earlier and increased SPCH expression, this however leads to an overall loss of patterning. Strikingly neither early ectopic expression of SPCH nor hyperactive SCRM-D, can force EPF2 or TMM expression or ACDs before the torpedo stage (Figure 3M-O, 6K-L).

A final developmental pause is before the completion of the stomatal lineage and formation of guard cells. We originally hypothesized, based on lack of embryonic expression, that FAMA was the limiting factor in embryonic stomatal formation. However, supplying embryonic FAMA (or elevating MUTE) still failed to induce stomata during embryogenesis (Figure 6E-G). Delayed differentiation of cell types specified during embryogenesis is not uncommon; for example, vascular tissues are specified and patterned but do not differentiate during embryogenesis (De Rybel et al., 2013; Truernit et al., 2008). What remain open questions, however, are how the pauses are generated, and how they are linked into to cell-type specific gene regulatory networks (GRNs). In the context of the stomatal GRN we see that both MUTE and FAMA can perform some of their regular functions during embryogenesis but cannot induce guard cell differentiation. Embryonic MUTE can only activate the cell division program during embryogenesis and FAMA can only induce a direct target gene, *ERF51*, and minor cell shape changes (Figure 6F-J, Figure 7A).

We hypothesize that prohibition of stomatal differentiation and the limited function of MUTE and FAMA during embryogenesis are linked to global mechanisms preventing differentiation during embryogenesis. Such mechanisms might include (1) a large overhaul in the transcriptome including unidentified stomatal cofactors linked to changes chromatin accessibility and DNA methylation (Narsai et al., 2017; van Zanten et al., 2011; Yamamuro et al., 2014), (2) differentiation of underlying tissue layers and resulting changes in cell-cell signaling (Baillie and Fleming, 2020; Wang et al., 2021) and/or (3) changes in global signaling resulting from hormone levels or exposure to light (Balcerowicz et al., 2014; Corbineau et al., 2014; Finkelstein et al., 2008; Lee et al., 2017). Preliminary studies using chemical inhibitors of chromatin modification and opening of siliques to expose developing ovules to light did not produce stomata in embryos (data not shown). Thus, further research is needed to determine whether and how a combination of these changes is needed for the progression of stomatal development. In addition, differences in MUTE and FAMA abilities between seedling and embryo could be explored to identify their unique transcriptional subroutines or co-factors.

### Embryonic stomatal patterning guides future patterning and allows for continuous rapid growth upon germination

Without a stomatal lineage (as in *spch* embryos) cells in the embryonic protoderm can still grow and divide, leaving open the question of whether it is essential for stomatal lineages to be initiated during embryogenesis. Our analysis of newly germinated seedlings that lack embryonic stomatal patterning, however, showed that all post-germination epidermal cells can then accumulate and respond to SPCH induction, resulting in ectopic lineage entry. Thus, pattern formation in a small surface where lineage specification and progression are slow, allows for gradual patterning and cell fate restriction in the absence of existing cues.

In addition, embryonically initiated stomata quickly differentiate upon germination. As non-embryonic stomata make up only ∼ 8% of total stomata in 2-day old wild type plants (Figure 4F), this suggests a major physiological role for embryonically patterned stomata in the period when the plant switches from relying on seed energy stores to fueling its own growth through photosynthesis. Early initiation of gas exchange for photosynthesis and resultant rapid growth are crucial for seedling establishment in field conditions (Finch-Savage and Bassel, 2016; Gommers and Monte, 2018).

### Slowing down *de novo* stomatal patterning allows for precision and informs the future pattern

The relatively delayed stomatal lineage entry contrasts with the establishment of other tissue identities and patterns during embryogenesis. Most cell identities and tissue patterns take shape as soon as the Arabidopsis embryo contains sufficient cells to form that particular identity or pattern and a variety of cell identities and patterns are already locked in at heart stage (ten Hove et al., 2015). Instead, stomatal lineage entry through asymmetric division is delayed until the bent cotyledon stage embryo, in contrast with lineage progression in the emerging true leaf where stomata quickly upon outgrowth (Figure S1H,L). The slow lineage patterning during embryogenesis allows us to visualize the cellular dynamics of *de novo* pattern formation.

*De novo* pattern generally involves cells either interpreting a morphogen gradient, like in the French Flag model (Vadde and Roeder, 2020; Wolpert, 1969), or self-organizing through reaction-diffusion (RD) mechanisms (Landge *et al*., 2020; Turing, 1952). Many epidermal models across animal systems employ RD models including patterning of shark denticles or avian feathers (Cooper et al., 2018; Painter *et al*., 2018). Such an RD-like model fits well with the expression patterns of SPCH and its direct targets whose expression domains all narrow over time. Indeed a RD-like method has been used to model stomatal pattern formation starting from a blank slate (Horst *et al*., 2015). However, here we find that the stomatal pattern develops during surface expansion and experiences slow-downs before lineage entry, hinting at slower, potentially different dynamics at the *de novo* formation of this pattern. Furthermore, we find that the *tmm* embryonic phenotype does not resemble that of post-embryonic leaves is not predicted by the current model (Figure 2I)(Horst *et al*., 2015), suggesting different dynamics in the stomatal regulatory network during initial pattern formation.

## Acknowledgements

MES was supported by a NWO Rubicon Postdoctoral fellowship (019.193EN.018), AV by an EMBO Postdoctoral fellowship (ALTF 707–2012), and AM by the Austrian Science Fund (J4019-B29). DCB is an investigator of the Howard Hughes Medical institute.

## Material and methods

### Key resources table

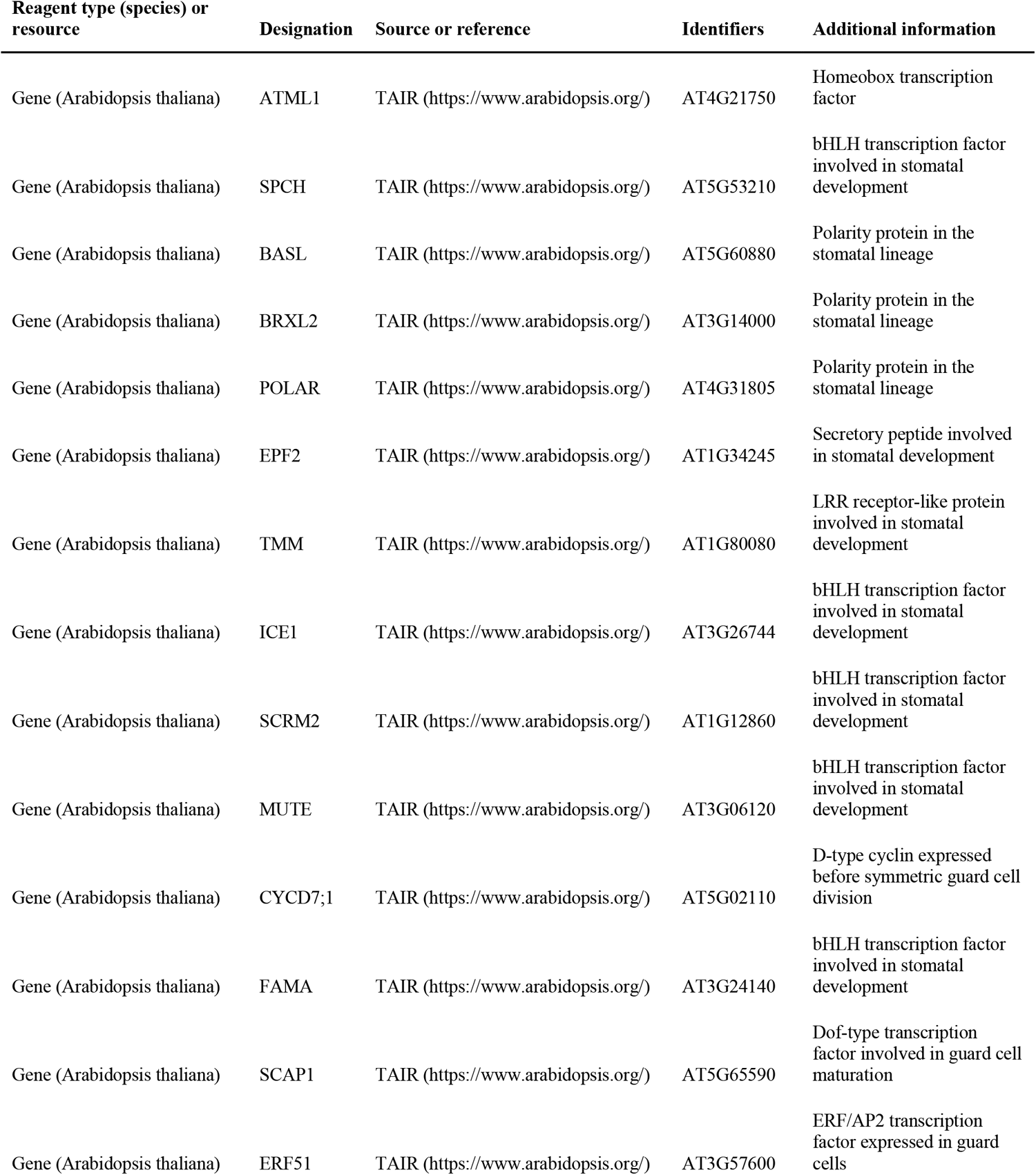
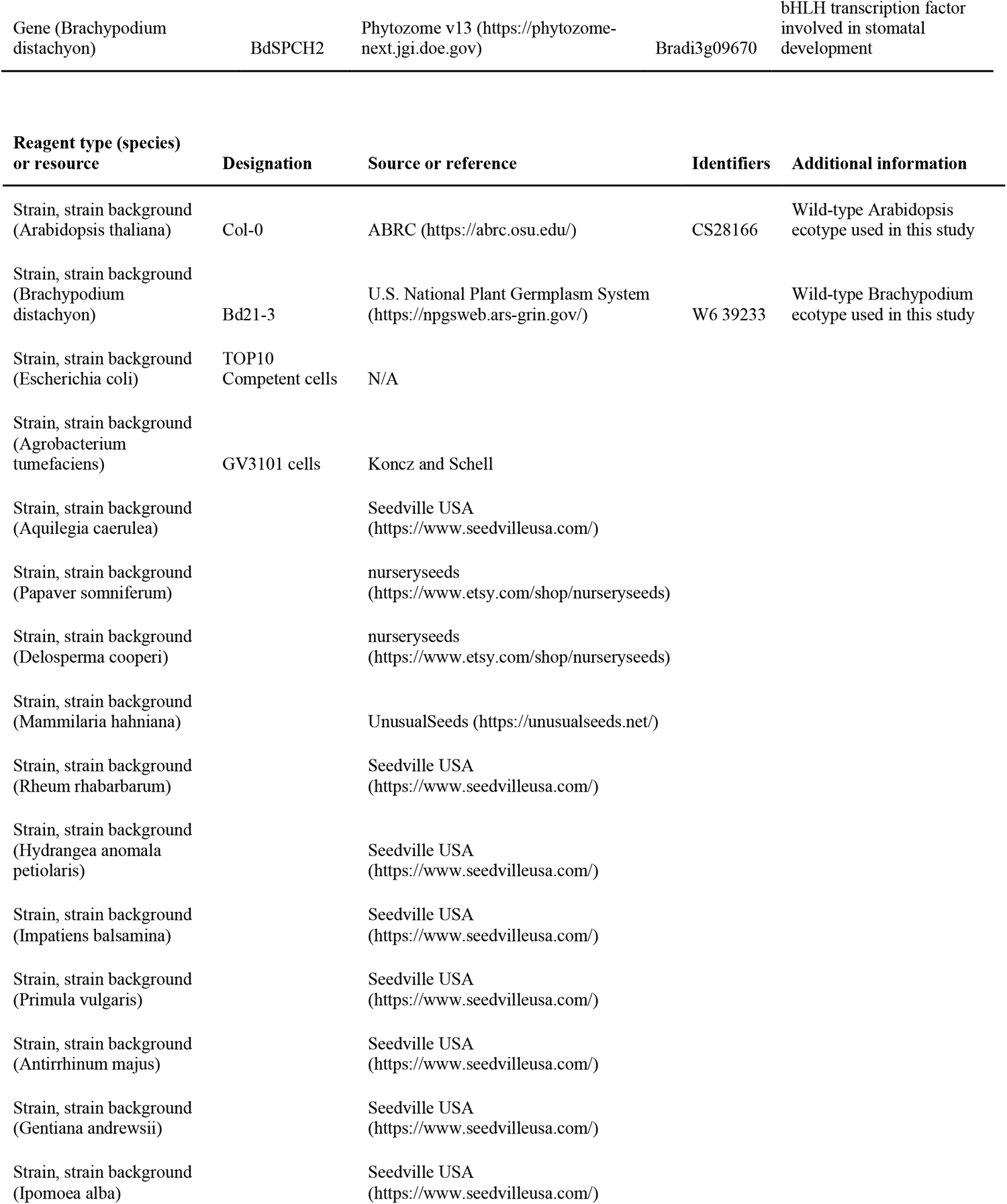
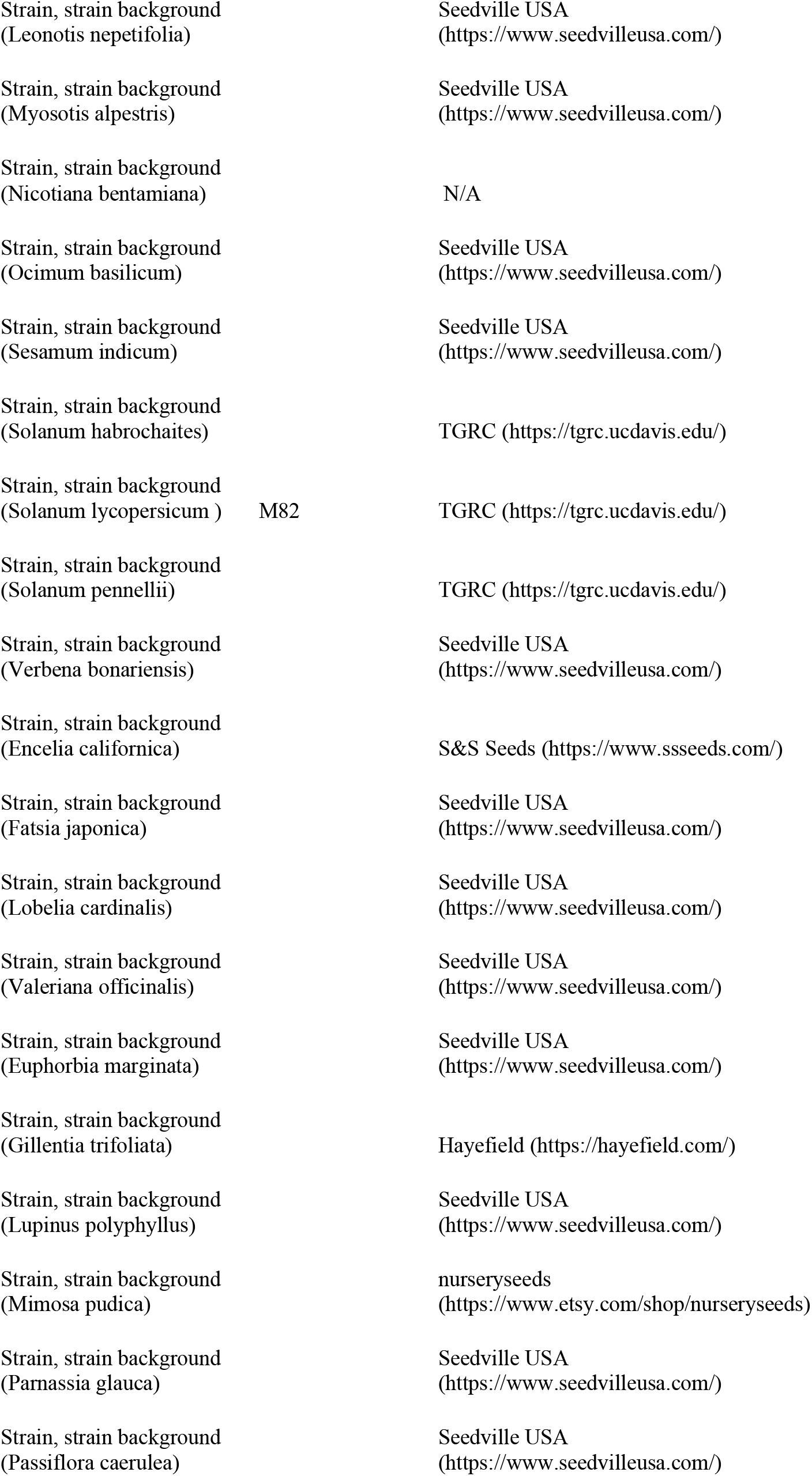
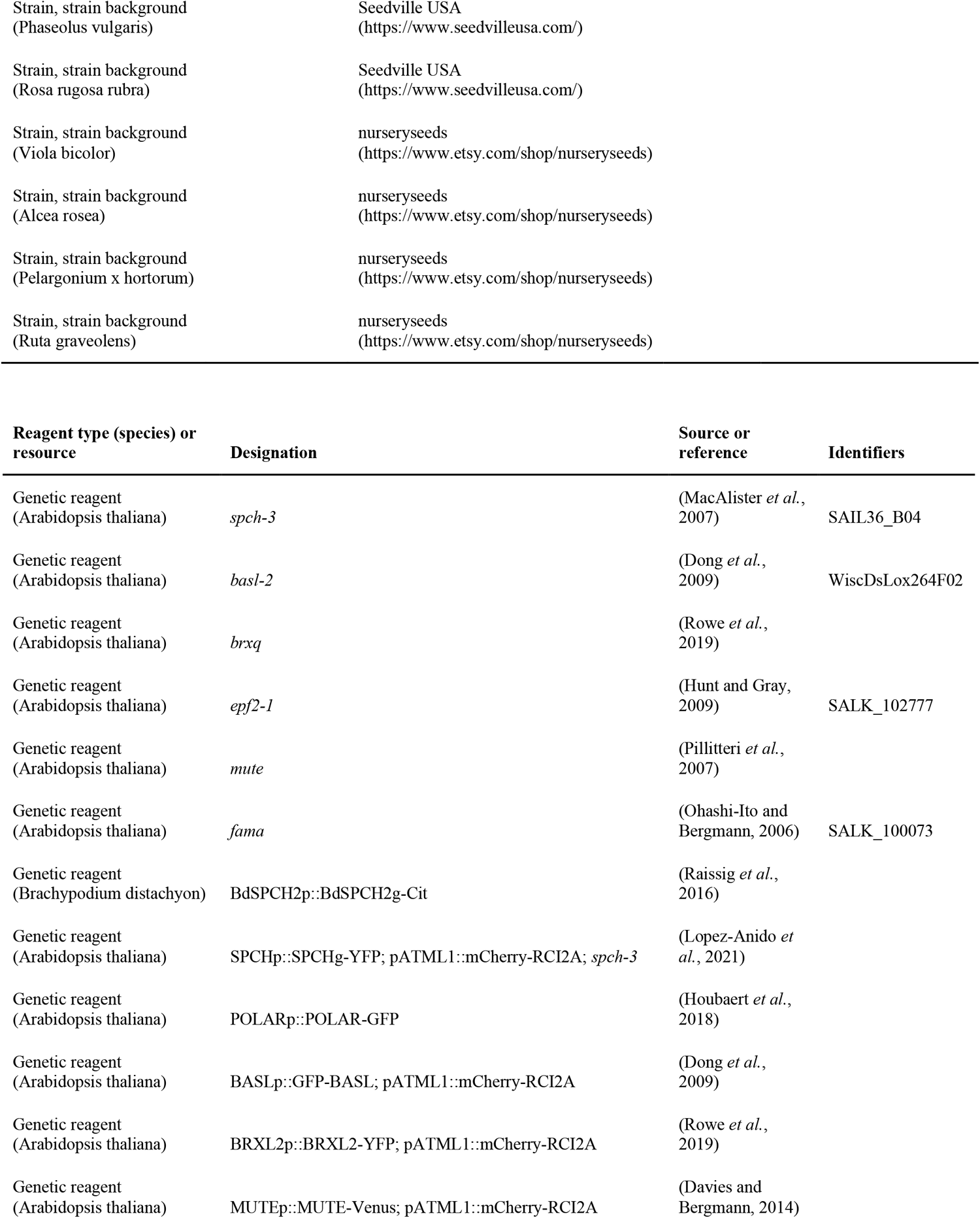
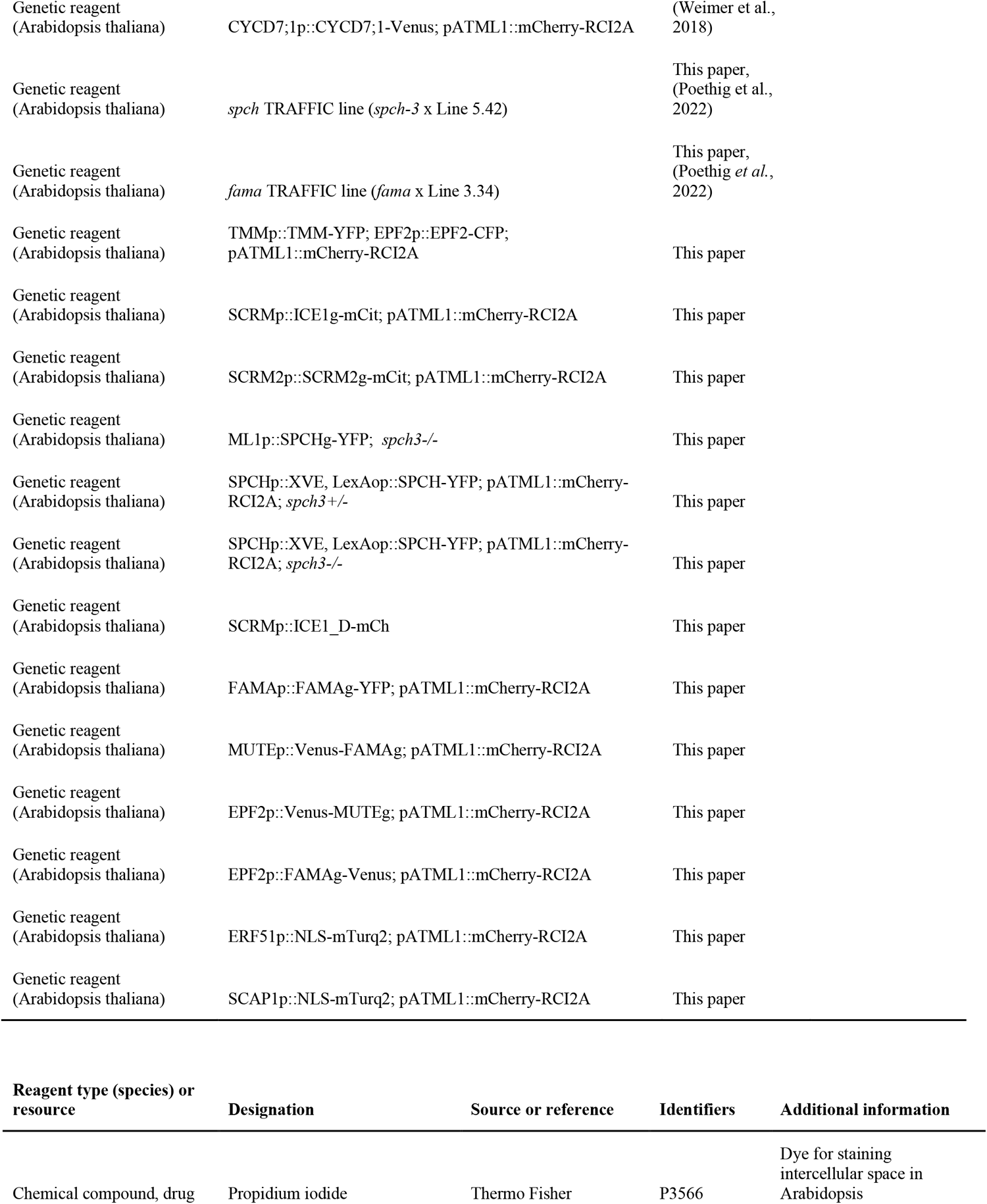
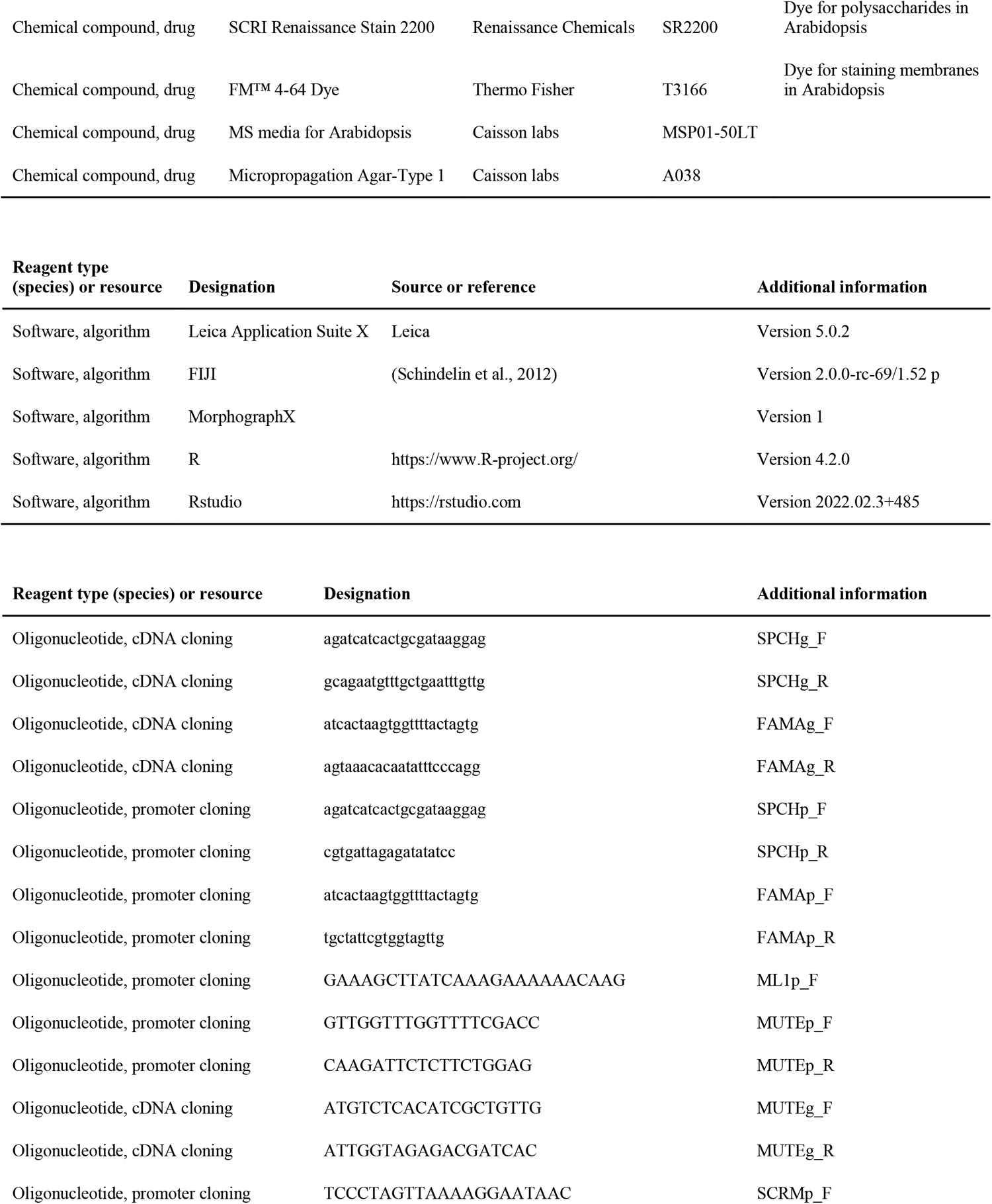
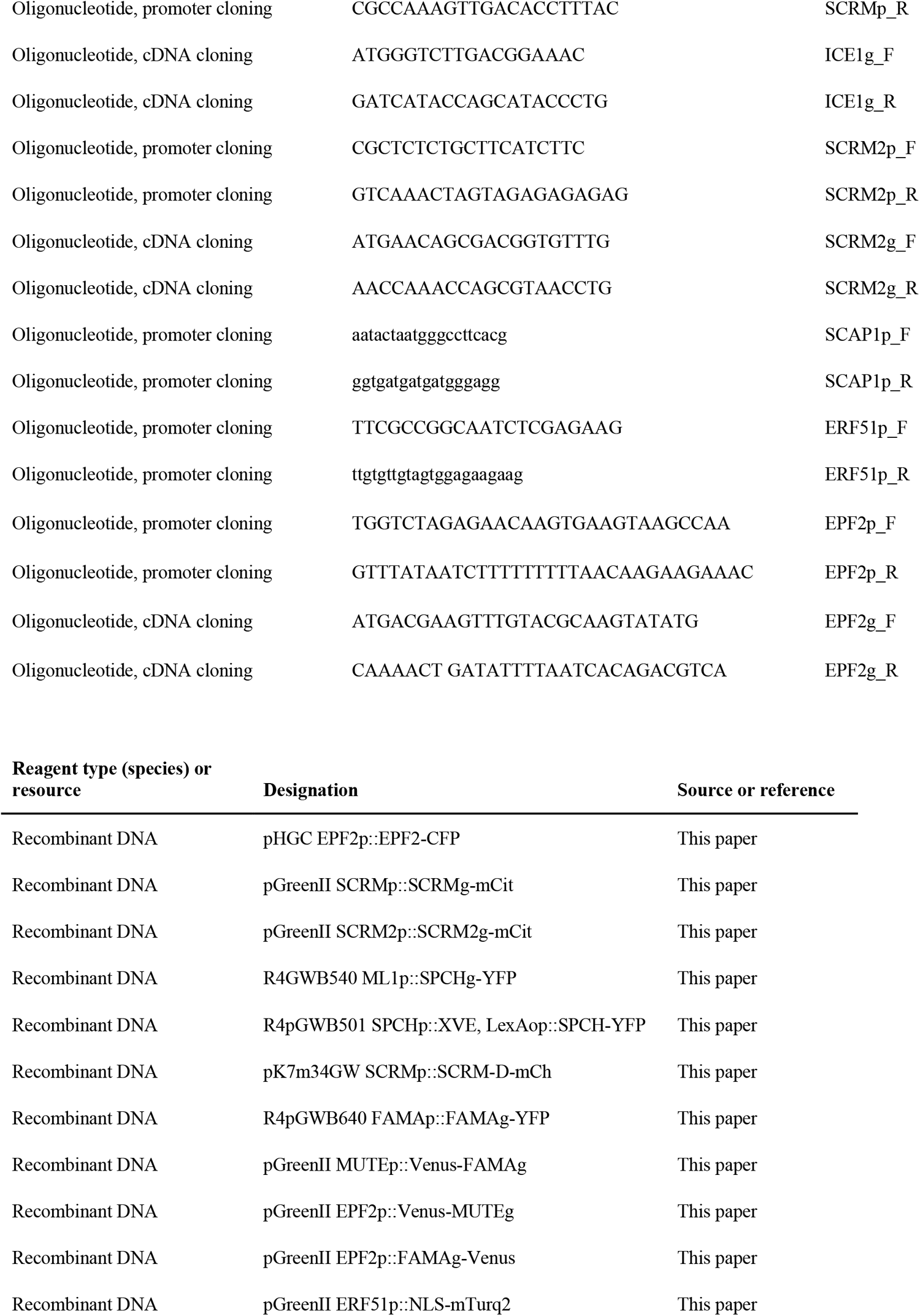
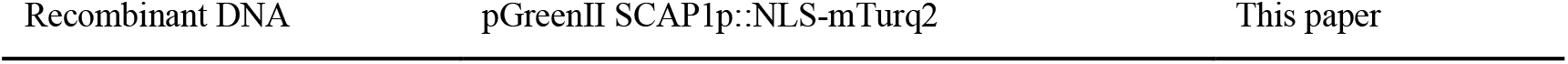

### Plant material and growth conditions

Arabidopsis lines used in this work are in the Col-0 background, newly generated lines and sources of previously reported transgenic lines are listed in the Key resources table. *spch, mute* or *fama* homozygous mutant seeds for dissection or timecourse experiments were generated or selected using inducible complementation (SPCHp::XVE, LexAop::SPCH-YFP; pATML1::mCherry-RCI2A in *spch-/*-) or TRAFFIC lines (for *spch, mute* and *fama*)(Poethig *et al*., 2022)(Key Resources table).

Arabidopsis thaliana seeds were surface-sterilized, plated on half-strength Murashige and Skoog medium without sucrose and with 0.8% agar. After 2 days of stratification at 4 °C, seedlings were grown for 2-14 days under standard long-day conditions in a Percival growth chamber, model CU36L5 (16 hr light/8 hr dark at 110 μmol m2 s-1 and 22°C).

Brachypodium distachyon Bd21-3 seeds were stratified in moist soil for 5 days at 4 °C before being moved to a Percival growth room at 26ºC with a 16-hour light/8-hour dark cycle (250 μmol m− 2 s− 1).

### Vector construction and plant transformation

Overview of all vectors created for this work in the Key Resources table. Constructs for plant transformation were cloned using GreenGate (Lampropoulos et al., 2013) backbone pGreenII or Gateway (Karimi et al., 2007) backbones pHGC, R4GWB540, R4pGWB501 or R4pGWB640. Transgenic plants were generated by floral dip (Clough and Bent, 1998) and transgenic seedlings were selected on ½ MS without sucrose with the appropriate antibiotic (15 mg/L phosphothricin, 7,5 mg sulfadiazine or 50 mg/L hygromycin).

### Sample preparation, microscopy and image processing

For consistency, abaxial leaf surfaces were imaged where possible. In developing embryos adaxial surfaces were imaged when accessible. In mature embryos the abaxial surface of the cotyledon facing away from the embryonic root was imaged. For larger embryos in the species panel cotyledons had to be cut up before mounting and it is unclear which leaf surface was imaged.

Stomatal reporters during embryogenesis were imaged using fresh embryos for Arabidopsis and after ClearSee treatment for Brachypodium embryos as described in (Hao *et al*., 2021).

Desiccated embryos were dissected from imbibed seeds and stained with mPS-PI and mounted with Hoyer’s (containing chloral hydrate)(Berleth and Jurgens, 1993; Truernit *et al*., 2008). Stacks were processed using MorphographX, creating a mesh to mark the tissue boundary onto which to project epidermal signal (2-8 μm from the mesh) before segmenting cell areas (de Reuille et al., 2015).

Stomatal lineage-like cells (Lineage Precursors, LPs) can be recognized by cell size and shape and tissue organization, time-course experiments and retrospective lineage tracing, and molecular markers (Gong et al., 2021; Nir et al., 2022). In the absence of molecular markers, such as in Figure 3, we relied on cell size and shape and on tissue context. For time-course (lineage tracing) experiments starting at radicle protrusion, germinating seeds were gently dissected 24-36 hours after moving to light, and the seedling was placed in water between to coverslips with vacuum grease providing mechanical support. This way both surfaces could be acquired without remounting the sample. After each time point, seedlings were carefully unmounted and placed on a ½ MS plate and normal growth condition until the next image acquisition. Lineage tracing was performed manually.

Fluorescence imaging experiments were performed on a Leica SP5, SP8 or Stellaris confocal microscope with HyD detectors using a 25x NA1.1 water objective. Cell outlines were visualized using a plasma membrane marker *ML1p::mCherry-RCl2A* (Davies and Bergmann, 2014) or one of the following dyes: propidium iodide (10 μ g/mL), FM4-64 (10 μM) or Renaissance 2200 (0.1%)(Musielak et al., 2015). For all stomatal reporter images, raw fluorescence image Z-stacks were projected with Sum Slices in FIJI unless otherwise stated.

### Induction experiments and analysis

To create embryos lacking SPCH, SPCHp::XVE, LexAop::SPCH-YFP; *spch3-/-* seedlings were germinated on 256 nM estradiol and after being transferred to soil were treated with estradiol every week until bolting. Upon bolting, estradiol treatment was halted. To select SPCHp::XVE, LexAop::SPCH-YFP; *spch3+/-* parents, seeds from a single parent were grown in the absence of estradiol and those with 3:1 survival were selected for experiments.

For phenotyping experiments, seeds were grown on 0 – 1024 nM estradiol for 4 days upon which whole leaf images were acquired. Images were stitched and projected to get a whole leaf view. Each image was then printed with name and treatment on the back. Pages were then shuffled, and phenotypes were scored blind.

## Notes

### Competing Interest Statement

The authors have declared no competing interest.

