## Supplemental Figures S1-5 for "Slow and not so furious: *de novo* stomatal pattern formation during plant embryogenesis"

Figure S1. SPCH expression during embryogenesis and first true leaf emergence

Figure S2. Increased ACD in ML1p::SPCH and SCRM-D embryos

Figure S3. Embryonic stomatal patterning is conserved across dicot species

Figure S4. Pre-patterned GMC-like cells rapidly give rise to stomata upon germination

Figure S5. Stomatal pre-patterning guides post embryonic stomatal development

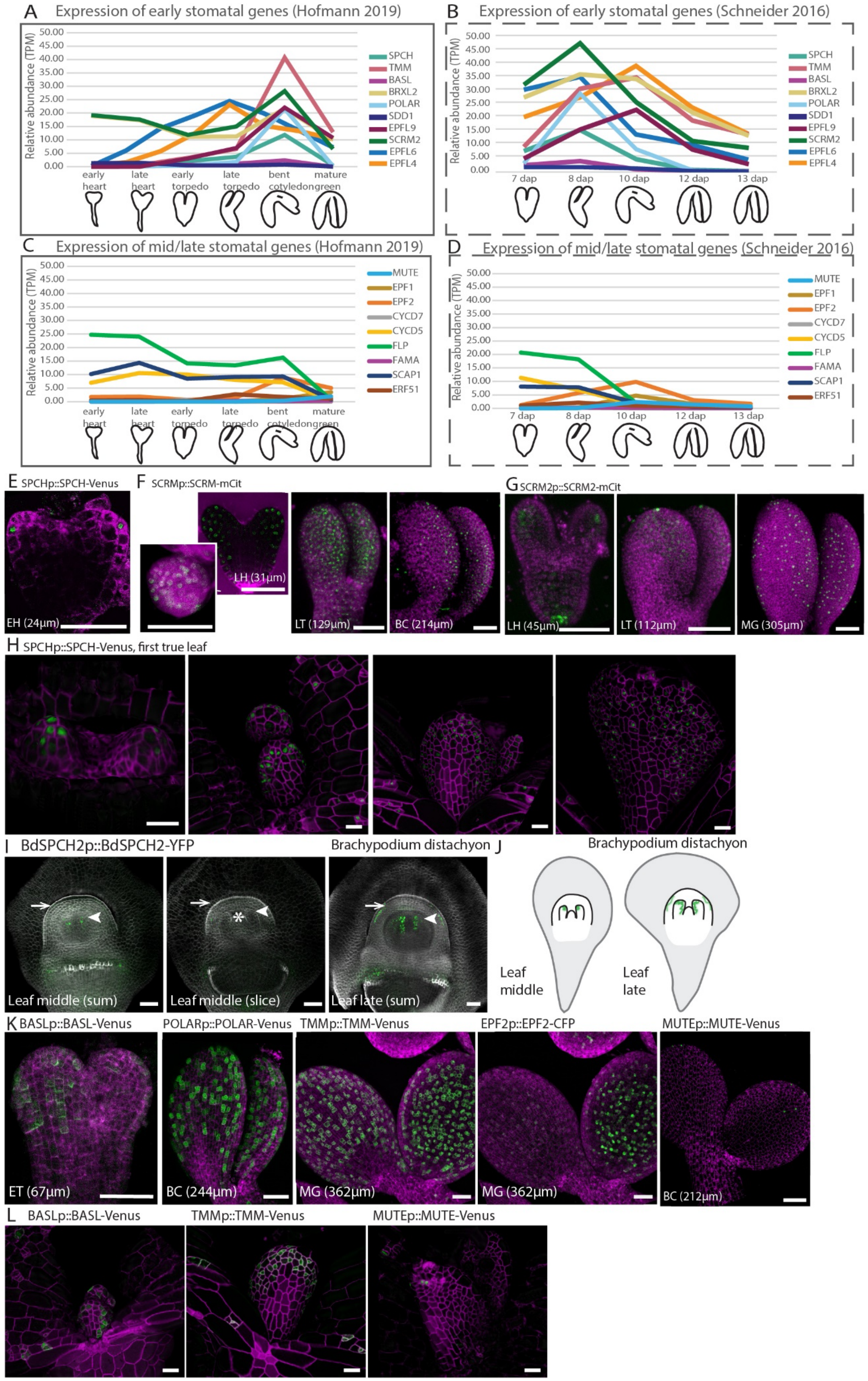

A Expression of early stomatal genes (Hofmann 2019)
B Expression of early stomatal genes (Schneider 2016)
C Expression of mid/late stomatal genes (Hofmann 2019)
D Expression of mid/late stomatal genes (Schneider 2016)
Relative abundance (TPM)
early heart, late heart, early torpedo, late torpedo, bent cotyledon, mature green
7 dap, 8 dap, 10 dap, 12 dap, 13 dap
SPCH, TMM, BASL, BRXL2, POLAR, SDD1, EPFL9, SCRM2, EPFL6, EPFL4
MUTE, EPF1, EPF2, CYCD7, CYCD5, FLP, FAMA, SCAP1, ERF51
E SPCHp::SPCH-Venus
EH (24μm)
F SCRMp::SCRM-mCit
LH (31μm)
LT (129μm)
BC (214μm)
G SCRM2p::SCRM2-mCit
LH (45μm)
LT (112μm)
MG (305μm)
H SPCHp::SPCH-Venus, first true leaf
I BdSPCH2p::BdSPCH2-YFP
Brachypodium distachyon
Leaf middle (sum)
Leaf middle (slice)
Leaf late (sum)
J Brachypodium distachyon
Leaf middle
Leaf late
K BASLp::BASL-Venus
POLARp::POLAR-Venus
TMMp::TMM-Venus
EPF2p::EPF2-CFP
MUTEp::MUTE-Venus
ET (67μm)
BC (244μm)
MG (362μm)
MG (362μm)
BC (212μm)
L BASLp::BASL-Venus
TMMp::TMM-Venus
MUTEp::MUTE-Venus

Figure S1. SPCH expression during embryogenesis and first true leaf emergence

**(**A-D) Transcript abundance of stomatal genes as reported in (Hofmann et al., 2019)(A,C) and (Schneider et al., 2016)(B,D). Split into early (A-B) and mid-late (C-D) stomatal genes. (E) SPCH expression (green) as reported by SPCHp::SPCH-YFP at early heart stage. (F-G) Expression of ICE1 (F) and SCRM2 (G) translational reporters during embryogenesis. Transition and late heart stage embryos are slices to show internal expression, later stages are stacks. (H) Expression of SPCH during emergence of the first true leaves. (I) Expression of BdSPCH2 during *Brachypodium distachyon* embryogenesis at leaf middle stage (left) and leaf late stage (right). Arrow points to the coleoptile, arrowhead points to the first leaf. Embryos were destained with Clearsee as described in methods. (J) Graphical summary of embryonic BdSPCH expression. (K) Additional images of SPCH target gene expression. (L) Expression of SPCH target genes and MUTE during emergence of the first true leaves. Magenta signal shows ML1p::mCherry-RCI2A and green signal a translational stomatal reporter. Scale bars represent 50 µm, tags indicate Embryo Stage (cotyledon length).

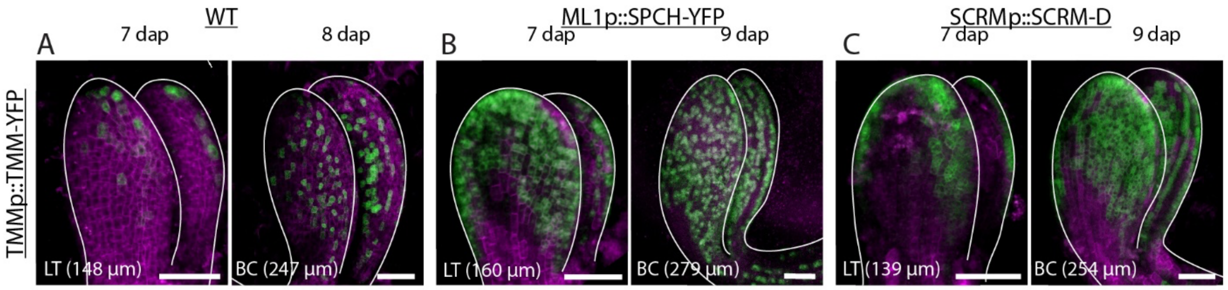


Figure S2. Increased ACD in ML1p::SPCH and SCRM-D embryos

(A-C) Combined membrane marker and TMM expression indicate increased ACDs in ML1p::SPCH-YFP (B) and SCRM-D (C) embryos compared to WT (A). Magenta signal shows ML1p::mCherry-RCI2A and green signal TMMp::TMM-YFP. Scale bars represent 50 µm, tags indicate Embryo Stage (cotyledon length).

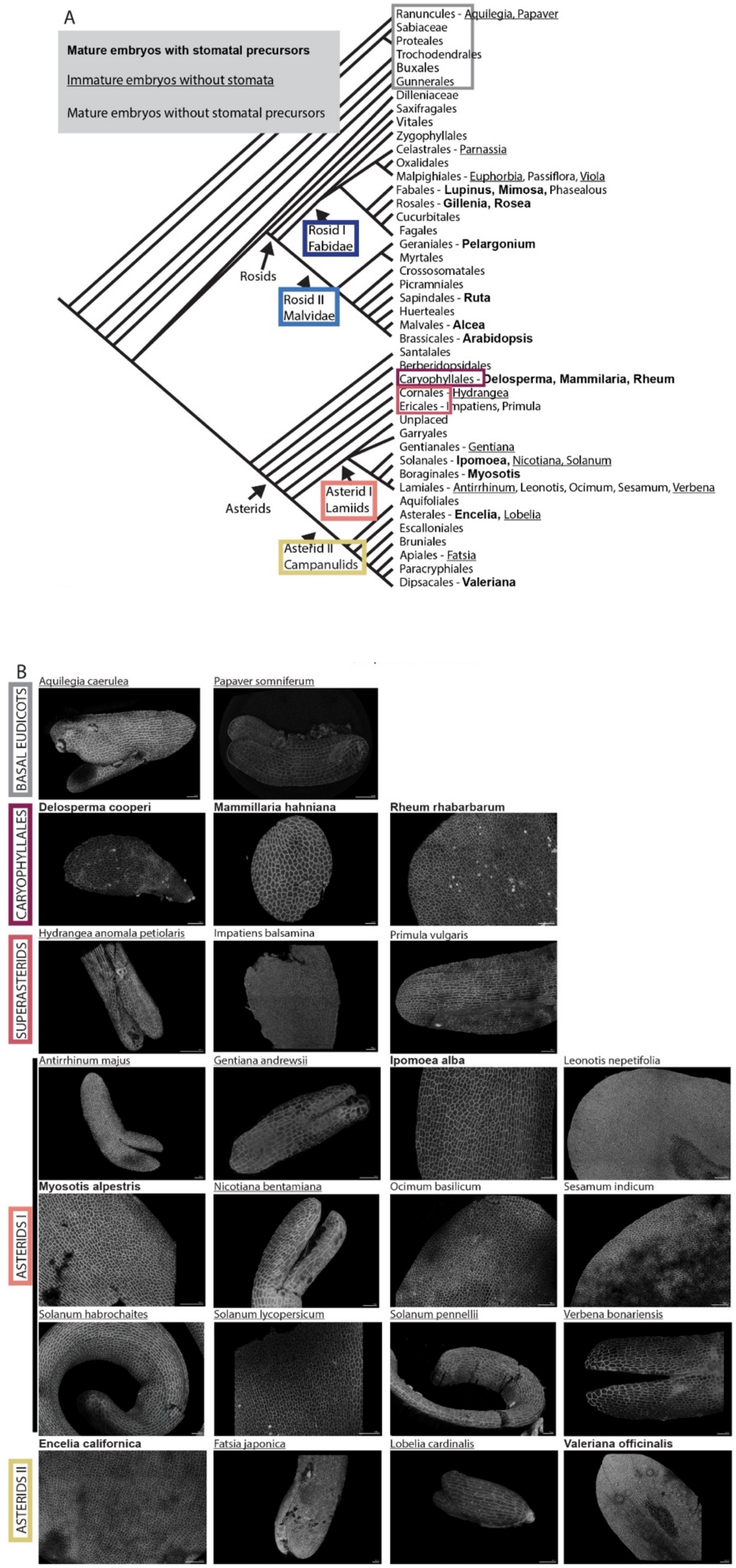
A
Mature embryos with stomatal precursors
Immature embryos without stomata
Mature embryos without stomatal precursors
Ranuncules - Aquilegia, Papaver
Sabiaceae
Proteales
Trochodendrales
Buxales
Gunnerales
Dilleniaceae
Saxifragales
Vitales
Zygophyllales
Celastrales - Parnassia
Oxalidales
Malpighiales - Euphorbia, Passiflora, Viola
Fabales - Lupinus, Mimosa, Phasealous
Rosales - Gillenia, Rosea
Cucurbitales
Fagales
Geraniales - Pelargonium
Myrtales
Crossosomatales
Picramniales
Sapindales - Ruta
Huerteales
Malvales - Alcea
Brassicales - Arabidopsis
Santalales
Berberidopsidales
Caryophyllales - Delosperma, Mammilaria, Rheum
Cornales - Hydrangea
Ericales - Impatiens, Primula
Unplaced
Garryales
Gentianales - Gentiana
Solanales - Ipomoea, Nicotiana, Solanum
Boraginales - Myosotis
Lamiales - Antirrhinum, Leonotis, Ocimum, Sesamum, Verbena
Aquifoliales
Asterales - Encelia, Lobelia
Escalloniales
Bruniales
Apiales - Fatsia
Paracryphiales
Dipsacales - Valeriana
Rosid I Fabidae
Rosids
Rosid II Malvidae
Asterid I Lamiids
Asterids
Asterid II Campanulids
B
BASAL EUDICOTS
Aquilegia caerulea
Papaver somniferum
CARYOPHYLLALES
Delosperma cooperi
Mammillaria hahniana
Rheum rhabarbarum
SUPERASTERIDS
Hydrangea anomala petiolaris
Impatiens balsamina
Primula vulgaris
ASTERIDS I
Antirrhinum majus
Gentiana andrewsii
Ipomoea alba
Leonotis nepetifolia
Myosotis alpestris
Nicotiana bentamiana
Ocimum basilicum
Sesamum indicum
Solanum habrochaites
Solanum lycopersicum
Solanum pennellii
Verbena bonariensis
ASTERIDS II
Encelia californica
Fatsia japonica
Lobelia cardinalis
Valeriana officinalis

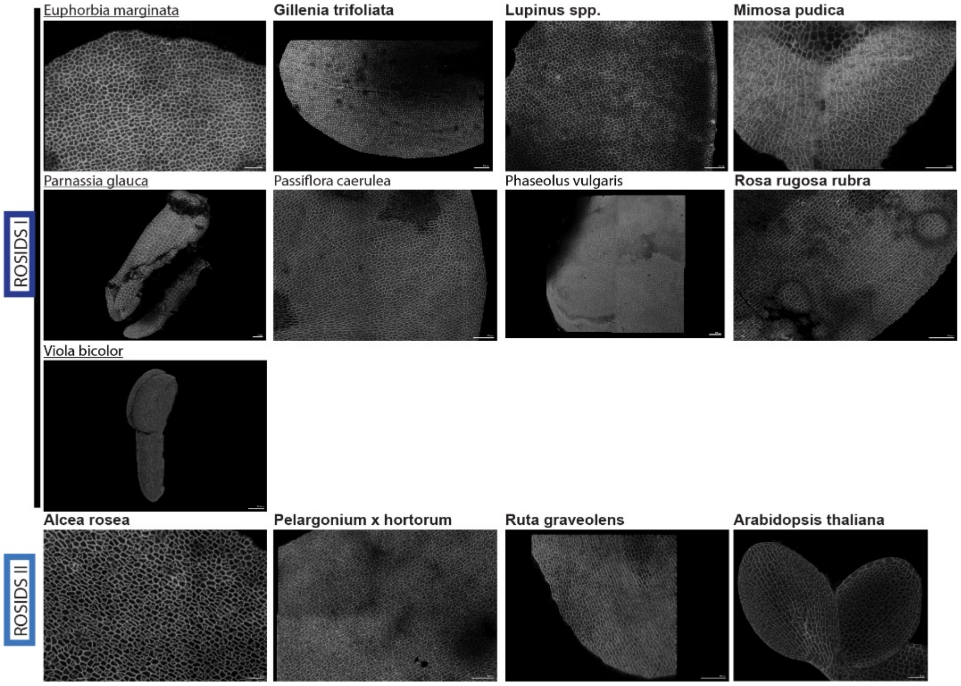


Figure S3. Embryonic stomatal patterning is conserved across dicot species
(A) Cladogram showing the families and specific plant species for which seeds were dissected. **Bold,** underline and regular indicate the presence of a stomatal pattern before germination and the size of the embryo (see legend). (B) Representative images of all species investigated. White signal shows propidium iodide signal after MorphographX processing (B). Scale bars represent 50 µm.

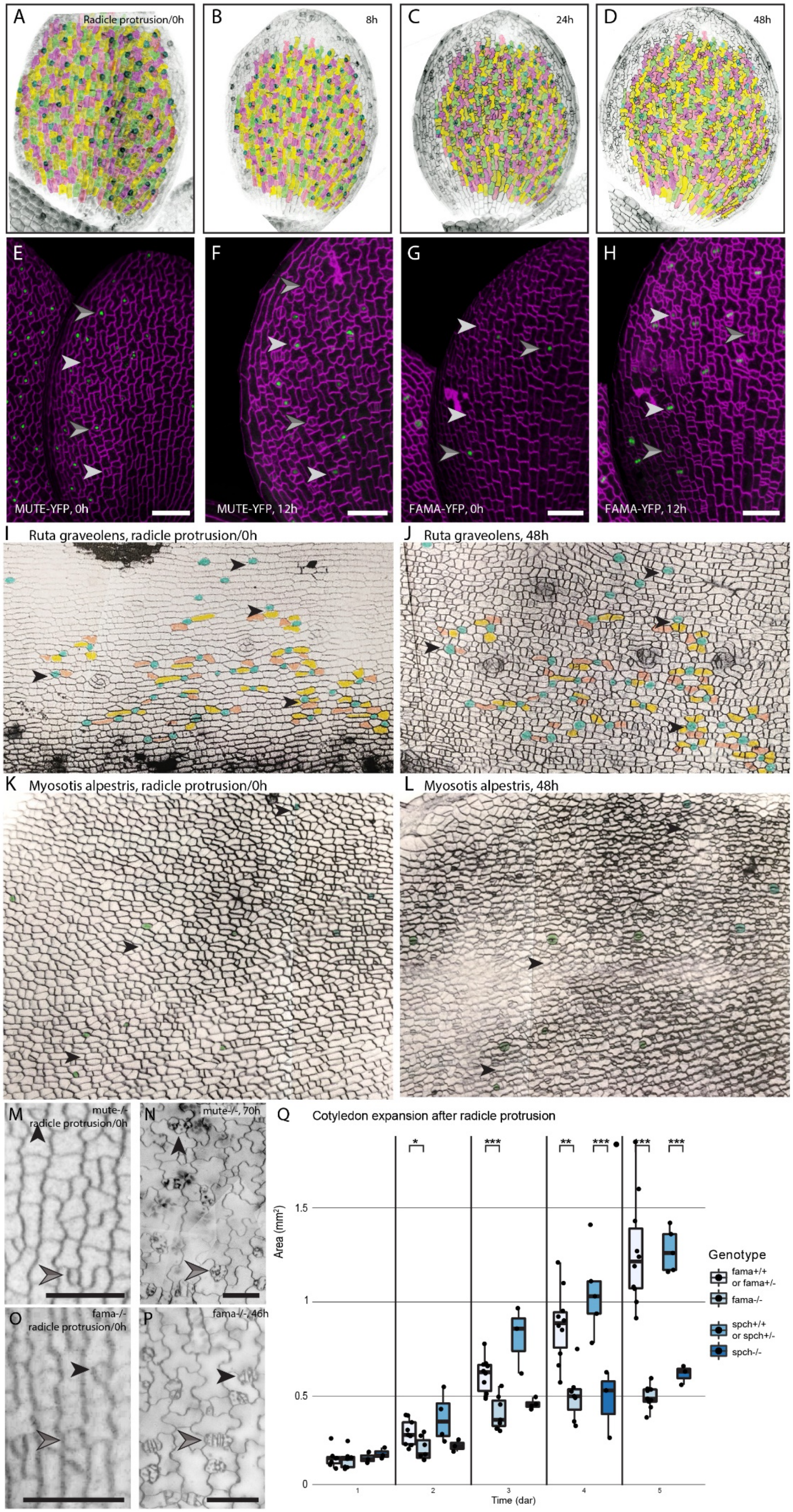

A
Radicle protrusion/0h
B
8h
C
24h
D
48h
E
MUTE-YFP, 0h
F
MUTE-YFP, 12h
G
FAMA-YFP, 0h
H
FAMA-YFP, 12h
I
Ruta graveolens, radicle protrusion/0h
J
Ruta graveolens, 48h
K
Myosotis alpestris, radicle protrusion/0h
L
Myosotis alpestris, 48h
M
mute-/-
radicle protrusion/0h
N
mute-/-, 70h
O
fama-/-
radicle protrusion/0h
P
fama-/-, 46h
Q
Cotyledon expansion after radicle protrusion
Area (mm2)
Time (dar)
Genotype
fama+/+ or fama+/-
fama-/-
spch+/+ or spch+/-
spch-/-

Figure S4. Pre-patterned GMC-like cells rapidly give rise to stomata upon germination.

(A-D) Time course series showing the same leaf as the main figure but with each cell lineage tracked to show its footprint until 48h after radicle protrusion. (E-H) MUTE (E-F) or FAMA (G-H) expression (green) at 0 and 12 hours after radicle protrusion. Grey arrowheads indicate cells which express MUTE/FAMA at 0h and which have divided within 12 hours, white arrowheads indicate LPs which don't express MUTE/FAMA at radicle protrusion, but which do express it 12 hours later. (I-L) Time course imaging of 2 non-model species (*Ruta graveolens, Myosotis alpestris*) whose embryonically patterned stomata cells directly differentiate, within 48 hours of radicle protrusion. (M-P) Time course series of *mute* (M-N) and *fama* (O-P) seedings at and several days after radicle protrusion. Stomatal cells look normal at radicle protusion but quickly diverge into characteristic division patterns. (Q) Cotyledon area of *fama* and *spch* seedlings at 1-5 days after radicle protrusion (dar), genotype was determined using seed coat fluorescence from TRAFFIC lines (see methods). *, ** and *** indicate p-values from pair-wise t-tests (<0.05, <0.01, <0.001). Black and magenta signal shows cell outlines with ML1p::mCherry-RCI2A or FM4-64, respectively. Scale bars represent 50 μm.

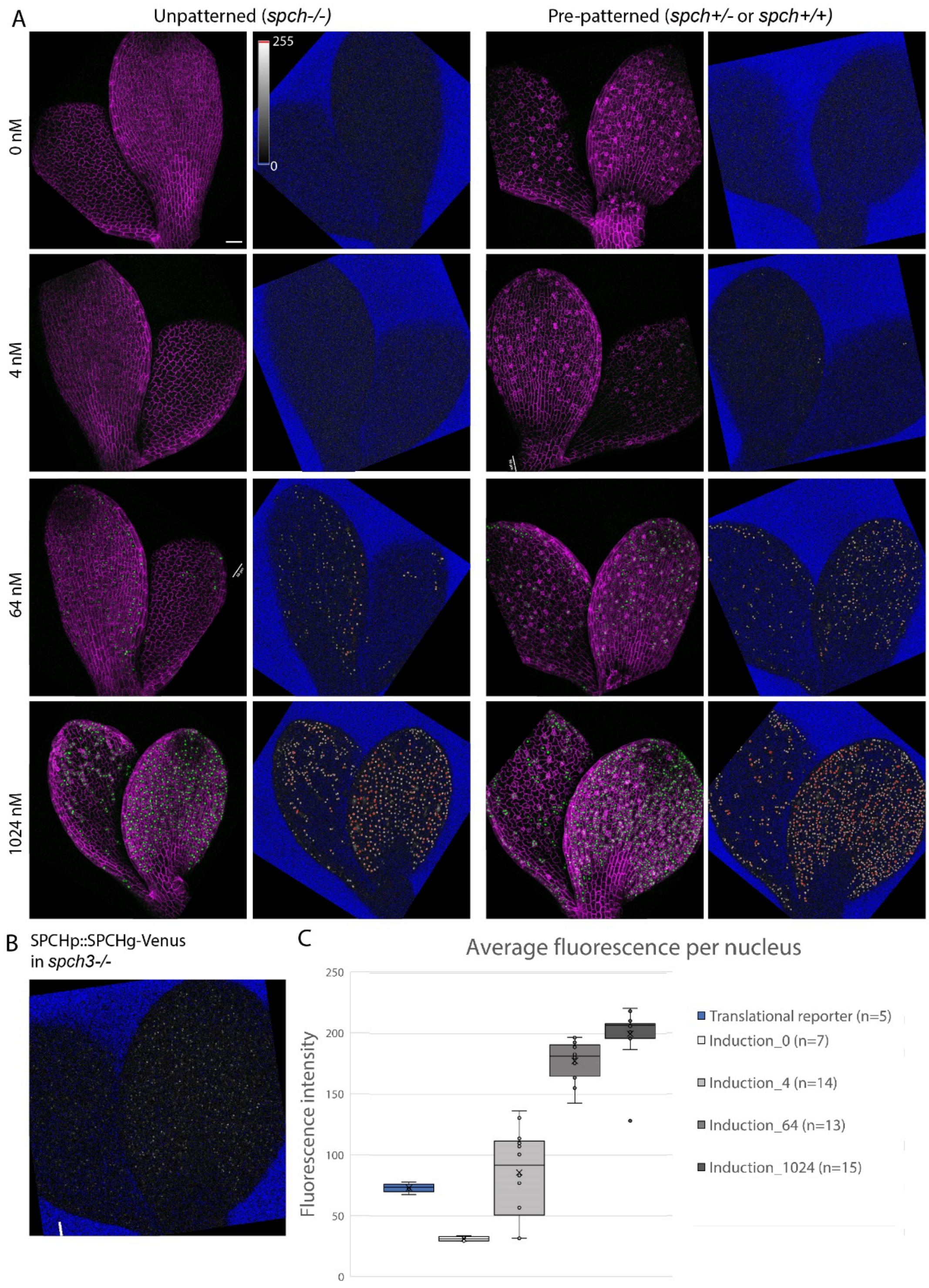


Figure S5. Stomatal pre-patterning guides post embryonic stomatal development.

(A) SPCH protein induction levels are similar between patterned and unpatterned cotyledons at 2 dag. Increasing estradiol concentrations resulted in both higher expression per cell and more cells expressing SPCH. Cell outline and SPCH (left) vs SPCH levels across the leaf (right). (B) SPCH protein levels in the complementation line. (C) Average fluorescence per nucleus across different levels of induction compared to a regular and strong SPCH complementation line. (A-B) SPCH localization (green) and membrane marker ML1p::mCherry-RCI2A (magenta) next to the YFP channel in the HiLo LUT. Scale bars represent 50 µm.
